# Diversification of small RNA pathways underlies germline RNAi incompetence in wild *C. elegans* strains

**DOI:** 10.1101/2021.08.21.457212

**Authors:** Han Ting Chou, Francisco Valencia, Jacqueline C. Alexander, Avery Davis Bell, Diptodip Deb, Daniel A. Pollard, Annalise B. Paaby

## Abstract

The discovery that experimental delivery of dsRNA can induce gene silencing at target genes revolutionized genetics research, by both uncovering essential biological processes and creating new tools for developmental geneticists. However, the efficacy of exogenous RNAi varies dramatically within the *C. elegans* population, raising questions about our understanding of RNAi in the lab relative to its activity and significance in nature. Here, we investigate why some wild-type strains fail to mount a robust RNAi response to germline targets. We observe diversity in mechanism: in some strains, the response is stochastic, either on or off among individuals, while in others the response is consistent but delayed. Increased activity of the Argonaute PPW-1, which is required for germline RNAi in the laboratory strain N2, rescues the response in some strains, but dampens it further in others. Among wild-type strains, genes known to mediate RNAi exhibited very high expression variation relative to other genes in the genome as well as allelic divergence and strain-specific instances of pseudogenization at the sequence level. Our results demonstrate functional diversification in the small RNA pathways in *C. elegans*, and suggest that RNAi processes are evolving rapidly and dynamically in nature.

## INTRODUCTION

In *C. elegans*, the landscape of molecular and developmental genetics was transformed by the ability to silence genes by feeding worms *E. coli* bacteria engineered to express RNA matching worm gene targets (*Kamath et al., 2003; Rual et al., 2004; Timmons & Fire, 1998*). This tool emerged from the seminal discovery that double stranded RNA induces gene suppression (*Fire et al., 1998*). Awarded the Nobel prize in 2006, the discovery launched research into new mechanisms of gene regulation and led to characterization of the microRNA (miRNA), short interfering RNA (siRNA), and PIWI-interacting (piRNA) pathways. These pathways overlap in gene set and molecular mechanisms, but mediate processes as diverse as cell growth and tissue differentiation, adaptive immunity against pathogens, transgenerational epigenetic inheritance, and germline defense against transposons (*Grishok, 2013; Wilson & Doudna, 2013*). Decades of investigation have illuminated a complex meta-phenomenon, with distinct sub-processes including the regulation of transcriptional silencing in the nucleus versus post-transcriptional silencing in the cytoplasm, of endogenous biogenesis of the small RNA trigger versus exogenous or environmental delivery, of local versus systemic responses, and of silencing genes in the soma versus targets in the germline. These processes are all captured under the umbrella term “RNA interference” (RNAi), the general mechanism of gene silencing via dsRNA (*Yigit et al., 2006*). Many of the genes that encode RNAi machinery are shared across plants, animals, and fungi, and appear deeply conserved within the eukaryotic lineage (*Shabalina & Koonin, 2008; Wynant et al., 2017*).

However, even as *C. elegans* sits at the epicenter of expansive research programs into gene silencing by small RNAs, wild-type strains vary significantly in capacity for RNAi. The universal laboratory strain N2 is robustly sensitive to RNAi, but its competence is not representative (*Félix, 2008*). For example, RNAi against germline targets in 55 wild isolates revealed a quantitative range of responses, from negligible to more sensitive than N2 *(Paaby et al., 2015*). Wild-type strains also vary in competence for targets in the soma, and some strains show incompetence for RNAi by both feeding and injection (*Félix et al., 2011; Paaby et al., 2015; Tijsterman et al., 2002*). To date, the only causal variant identified for natural differences in RNAi is a frameshift lesion in the Argonaute *ppw-1*, which partially explains germline RNAi insensitivity in the Hawaiian isolate CB4856 (*Tijsterman et al., 2002*). The genetics underlying differences in RNAi efficacy in *C. elegans* are otherwise unknown.

RNAi silences complementary mRNA targets via the association of small RNAs with Argonaute effector proteins (*Wilson & Doudna, 2013*). The Argonaute superfamily includes the ancient AGO proteins; the PIWI Argonautes, which are conserved in animals; and in *C. elegans*, the WAGO proteins, an expanded clade of Argonautes specific to nematodes (*Youngman & Claycomb, 2014*). The expansion of Argonautes signifies a critical role for RNAi in nematodes, and has been hypothesized to underlie the extraordinary diversification of worms across environments, perhaps having enabled adaptations associated with environmental sensing, parasitism, and immunity (*Buck & Blaxter, 2013*). Likewise, the momentum of discovery of RNAi-associated phenomena at the bench increasingly indicates that gene regulation by small RNAs dominates *C. elegans* biology (*Houri-Zeevi et al., 2020; Youngman & Claycomb, 2014*).

Given the evident importance of RNAi in *C. elegans*, why is its efficacy so variable? Here, we investigate the genetic basis of germline RNAi deficiency in wild *C. elegans* strains. We aim to elucidate how the response fails in incompetent strains, the role of *ppw-1*, and whether the genetic architecture of incompetence is simple or complex. We also evaluate expression and allelic diversity at genes known to mediate RNAi, to compare RNAi responses at the organismal level to proximate causes of failure. These analyses uncover evidence of extensive diversification of RNAi activity within *C. elegans*, consistent with rapid and recent evolution of a genetically complex trait. This level of functional variability in RNAi pathways offers a useful access point into connecting the vast body of *C. elegans* RNAi research to the biological relevance of these processes in nature.

## RESULTS

### Germline RNAi varies in expressivity and penetrance over reproductive age and among genotypes

Prior work examining embryonic gene knockdown in wild *C. elegans* demonstrated that strains vary quantitatively in the strength of their germline RNAi response, and that strains CB4856 and QX1211 appear largely incompetent for germline RNAi (*Paaby et al., 2015*). In contrast, the common wild-type laboratory strain N2 is highly sensitive to RNAi, though germline RNAi can be eliminated in N2 with a deletion at the WAGO Argonaute *ppw-1* (*Tijsterman et al., 2002; Yigit et al., 2006*). To directly compare RNAi incompetence in the N2 mutant (N2*^ppw-1(del)^*), CB4856, and QX1211, we targeted the maternal-effect, embryonic-required genes *par-1* and *pos-1*, which have commonly been used to measure germline RNAi (*Elvin et al., 2011; Pollard & Rockman, 2013; Tijsterman et al., 2002*). We fed worms *E. coli* expressing target dsRNA, then counted dead embryos in the next generation. Under a conventional approach of pooling worms on plates and scoring offspring in a window of relatively early egg-laying (*Kamath et al., 2001; Pollard & Rockman, 2013*), our observations were consistent with prior reports: wild-type N2 exhibited high lethality, and the three incompetent strains exhibited very low or negligible lethality (Figure 1A).

**Figure 1.**
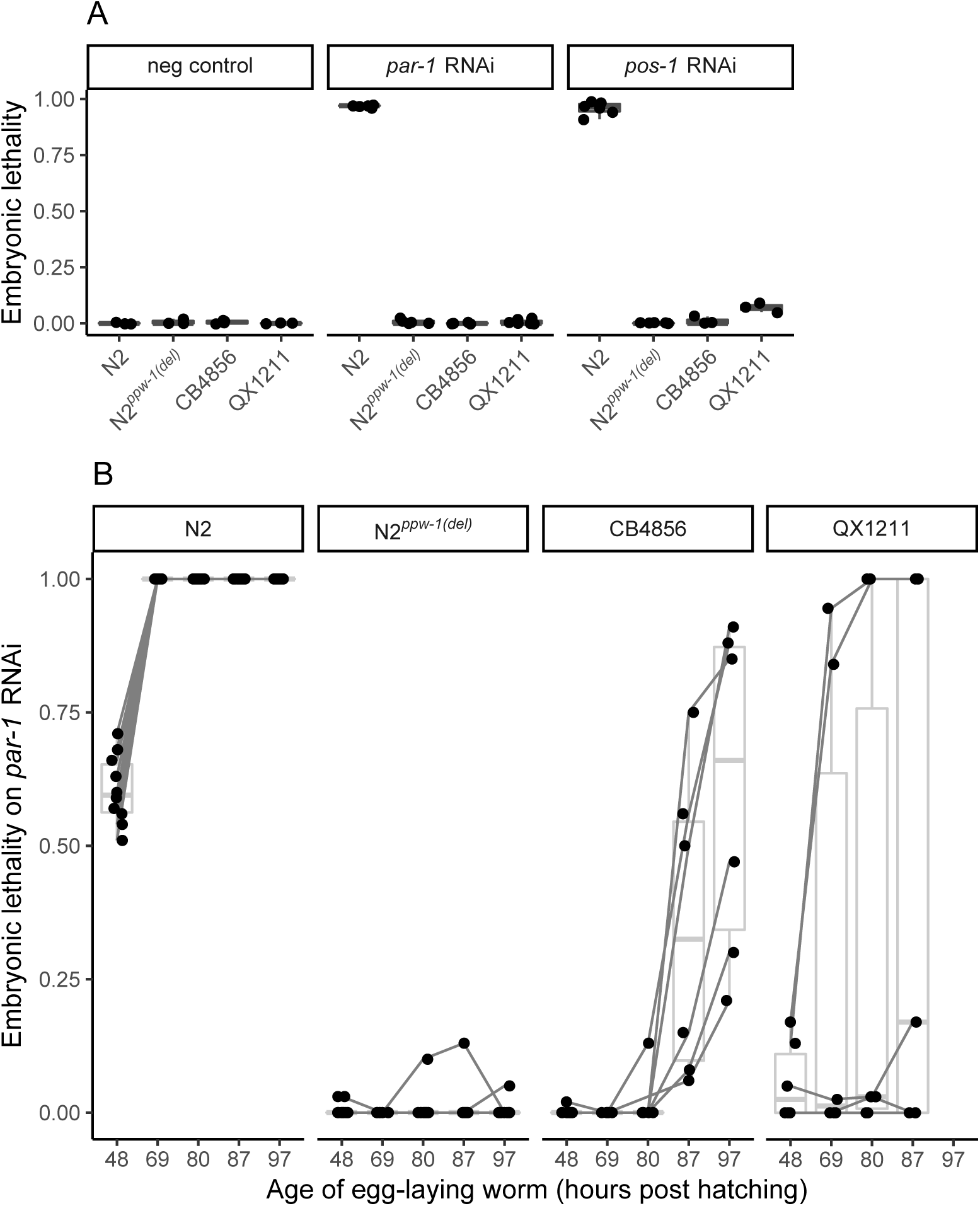
Embryonic lethality following RNAi against germline-expressed targets. (A) Hatched larvae and dead embryos laid in a 4-6hr window within the first 8hrs of egg-laying were scored for pooled hermaphrodites across replicate plates. (B) To assess the germline RNAi response over reproductive lifespan, embryonic lethality was scored for individuals. Each point represents the proportion of dead embryos, out of total laid on a plate by a single hermaphrodite, in the given time interval. The data include all offspring of all hermaphrodite mothers; time intervals were chosen to space out the number of offspring per plate (∼30-100); x-axis labels indicate the approximate midpoint of the time intervals. Connecting lines indicate the sequence of offspring plates for each individual. Embryonic lethality for all strains on the negative control empty vector was negligible (data not shown).

However, following an initial trigger, amplification within the RNAi response can induce strong effects later (*Billi et al., 2014*), and continual exposure to RNAi by feeding also means that individuals ingest increasing numbers of trigger molecules as they age. To evaluate whether the response changes over time within individual animals, we scored the penetrance of embryonic lethality over the complete reproductive lifespan of egg-laying individuals. In this assay we targeted *par-1*, which provides the more sensitive readout since it is not as lethal. Here, each of the three incompetent strains exhibited a distinct response, indicating differences in genetic mechanism (Figure 1B).

N2 showed complete lethality in all but the earliest offspring, suggesting that in this sensitive strain, early amplification of the initial trigger rapidly induces total gene knockdown. In the mutant N2*^ppw-1(del)^*, however, nearly all embryos hatched, including late-age embryos (Figure 1B), indicating that the loss of *ppw-1* is not compensated by other genes in the N2 background.

In CB4856, hermaphrodite mothers exhibited no evidence of an RNAi response in the first half of their reproductive lifespan, but embryonic lethality emerged in the second half and increased with parental age (Figure 1B). This suggests that the mutation in *ppw-1*—which encodes a frameshift and early stop upstream of the critical PAZ and PIWI domains and is putatively responsible for insensitivity in CB4856 (*Elvin et al., 2011; Pollard & Rockman, 2013; Tijsterman et al., 2002*)—is either not a null allele and permits some PPW-1 activity, or that other genes in the

CB4856 background partially compensate for the loss of PPW-1, promoting a delayed RNAi response.

QX1211 exhibited a third unique non-competent response. After a short delay, embryonic lethality was either negligible or complete, suggesting that RNAi in QX1211 is either “on” or “off” in individual animals (Figure 1B). Thus, unlike N2*^ppw-1(del)^*, in which the RNAi response appears abolished, CB4856 and QX1211 do exhibit limited responses, but with distinct patterns of activity: in CB4856, the response is delayed and incomplete; in QX1211, it is partially delayed, with higher expressivity and variable penetrance.

These results point to distinct differences in the execution of germline RNAi within *C. elegans*. However, the use of an end-point phenotype to read out the RNAi response, i.e. embryonic lethality, does not capture activity at the molecular or cellular level. Moreover, variation in the *par-1* pathway between strains might influence phenotypic expression, confounding interpretation of the RNAi response (*Paaby et al., 2015*). Therefore, we developed an assay to measure the expression and knockdown of the target gene directly.

### Target transcript knockdown confirms distinct RNAi responses across strains

To assess the RNAi response with high spatial and temporal resolution within embryos, we used single-molecule fluorescence *in situ* hybridization (smFISH) to visualize transcripts of germline RNAi targets. Since smFISH visualizes individual molecules via hybridization of dozens of oligonucleotide probes, which in aggregate produce a detectable fluorescent spot (*Raj et al., 2008*), it captures signals from intact RNAs, not those degraded by RNAi. These experiments confirmed that the distinct responses of CB4856 and QX1211 are driven by variation in RNAi mechanism, not in developmental variation related to the RNAi target. They also illustrated consistent transcript degradation across the early stages of embryogenesis.

We examined *par-1* transcript levels in *par-1* RNAi-treated and -untreated embryos of N2, CB4856 and QX1211. We collected embryos from gravid worms in early reproductive maturity, in a narrow two-hour window, to maximize precision in estimating the RNAi response. At this timepoint, many *par-1* transcripts are degraded in RNAi-treated N2 embryos (Figure 2A-B).

**Figure 2.**
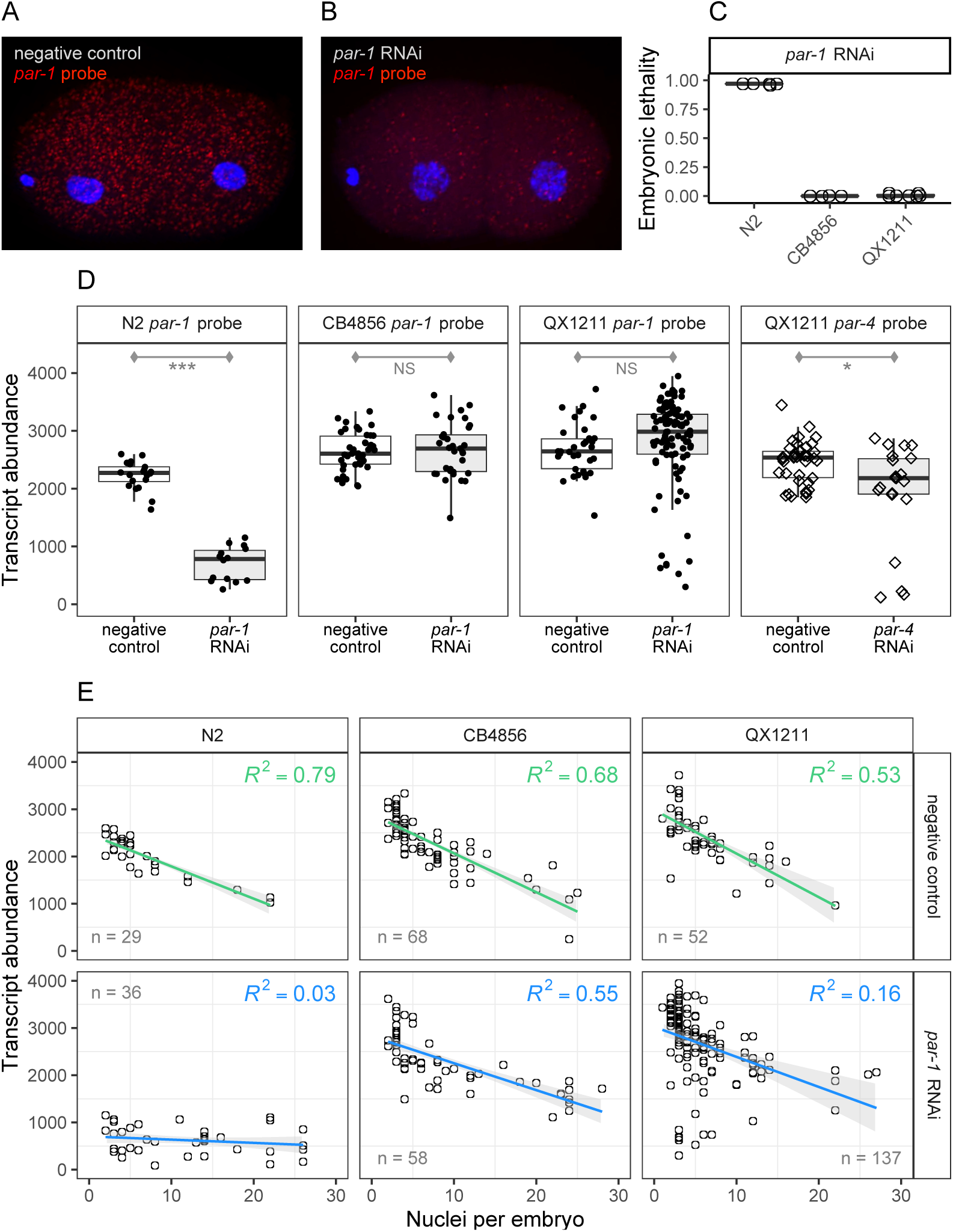
Transcript abundance in individual RNAi-treated and untreated embryos, visualized via single-molecule fluorescence in situ hybridization (smFISH). (A) - (B) Representative embryos are shown from strain N2, from a mother reared in the control condition or with RNAi against *par-1*; red spots indicate *par-1* transcripts and blue DAPI staining shows nuclei, which were used to identify embryo stage. (C) Embryonic lethality was simultaneously measured in matched samples. To limit variation due to reproductive age of the mothers, we collected embryos from a tightly controlled time window in early reproduction. (D) Transcript abundance for early stage embryos (up to four cells). (E) Transcript abundance for *par-1* for the same experiment, but now including later stage embryos with up to 30 nuclei. Green (negative control) and blue (RNAi treatment) lines indicate the linear regression of transcript counts onto embryonic stage; gray shading indicates the 90% confidence interval. For (D) and (E), each point represents one embryo. Significance levels (*t*-tests): p<0.001 (***), p<0.01 (**), p<0.05 (*).

Treated N2 embryos of this timepoint go on to show complete lethality, but in CB4856 and QX1211, lethality is not yet penetrant (Figure 2C).

All three strains displayed robust expression of the target gene in untreated embryos (Figure 2D), indicating that levels of native gene expression are unlikely to be a major influence on lethality penetrance. However, in RNAi-treated embryos, N2 showed a steep drop in transcript abundance, CB4856 showed no change, and QX1211 showed an on/off pattern with N2-like levels for some, but not most, embryos. This pattern in QX1211 was replicated for a second target, *par-4* (Figure 2D); see File S1 for statistical details. Thus, the patterns of transcript knockdown following RNAi are highly consistent with our prior observations of strain-specific responses.

To examine how transcript abundance, with and without degradation by RNAi, changes with embryonic development, we evaluated embryos with up to 30 nuclei. In the control condition, *par-1* transcripts decreased with embryonic stage (Figure 2E) (*Charles et al., 2021*) at a consistent rate across strains (ANCOVA model comparison, p=0.299), indicating no apparent differences in *par-1* developmental activity. In the treatment condition, the strain-specific patterns of transcript degradation persisted without any apparent effect of development on the RNAi response (Figure 2E). That is, the treated N2 embryos, following a significant knockdown in transcript number (p<0.001), exhibited a flat slope that implies no change in RNAi response with embryo stage. The treated CB4856 and QX1211 embryos exhibited negligible change by embryo stage relative to the control condition, with marginal (ω^2^=0.017, p=0.012) and non-significant changes in slope, respectively. (The complete statistical report for this analysis, including estimates of the variance explained and significance levels for ANCOVA model comparisons, is in Table S1.) Thus, in this narrow window of embryogenesis and among embryos retrieved from a fixed-age parent, we find no evidence of changing rate of degradation by embryo stage.

### Reduced PPW-1 function does not universally explain loss of germline RNAi

Given the distinct patterns of germline RNAi incompetence in CB4856 and QX1211, we next sought to evaluate the genetic basis for RNAi failure in these and other low-response strains. First, we first considered the role of *ppw-1*. The naturally-occurring frameshift mutation in *ppw-1* (*Tijsterman et al., 2002*) is unique to CB4856 in the *Caenorhabditis elegans* Natural Diversity Resource (CeNDR) database (*Cook et al., 2017*), but we hypothesized that variation in PPW-1 activity arising from other sources might contribute to variation in germline RNAi among wild strains. To test whether reduction of PPW-1 function is a universal aspect of reduced germline RNAi, we performed complementation tests by crossing N2 wild-type and null alleles of *ppw-1* to seven wild isolates. We evaluated CB4856, QX1211, and five additional strains, selected based on prior observations of weak germline RNAi (data not shown) (*Paaby et al., 2015*) and representation of nucleotide diversity and divergence across the global population (*Cook et al., 2017*).

We crossed each wild isolate to N2 with its native, wild-type copy of *ppw-1*, and also to N2 carrying the *ppw-1* deletion allele (*pk1425*). Two genetic incompatibilities segregating within *C. elegans* (*Ben-David et al., 2017; Seidel et al., 2008,* 2011) complicated our crosses, one of which we controlled with a knockout allele at *peel-1*; details are provided in File S2. For each cross, we compared the response of the individual wild isolate to the two heterozygote genotypes in the F1 generation, with and without the deletion at *ppw-1* inherited from the N2 chromosome. We reasoned: though N2 homozygous for the *ppw-1* deletion fails to exhibit an RNAi response, one copy of wild-type *ppw-1* fully rescues it, indicating that *ppw-1* is haplo-sufficient, at least in the N2 background (Figure 3A). Therefore, if weak RNAi in the wild strains is a consequence of reduced PPW-1 activity, any restoration of response in the F1 genotypes should be greater in the genotype with the functional N2 *ppw-1* allele. As previously, we induced *par-1* RNAi in the (F1) parent germline and measured embryonic lethality in the following generation. To avoid confounding differences in developmental timing with variation in RNAi response, we scored all progeny from only the first 15hrs of egg-laying from a small pool of hermaphrodite parents (∼100-200 embryos) on each replicate plate.

**Figure 3.**
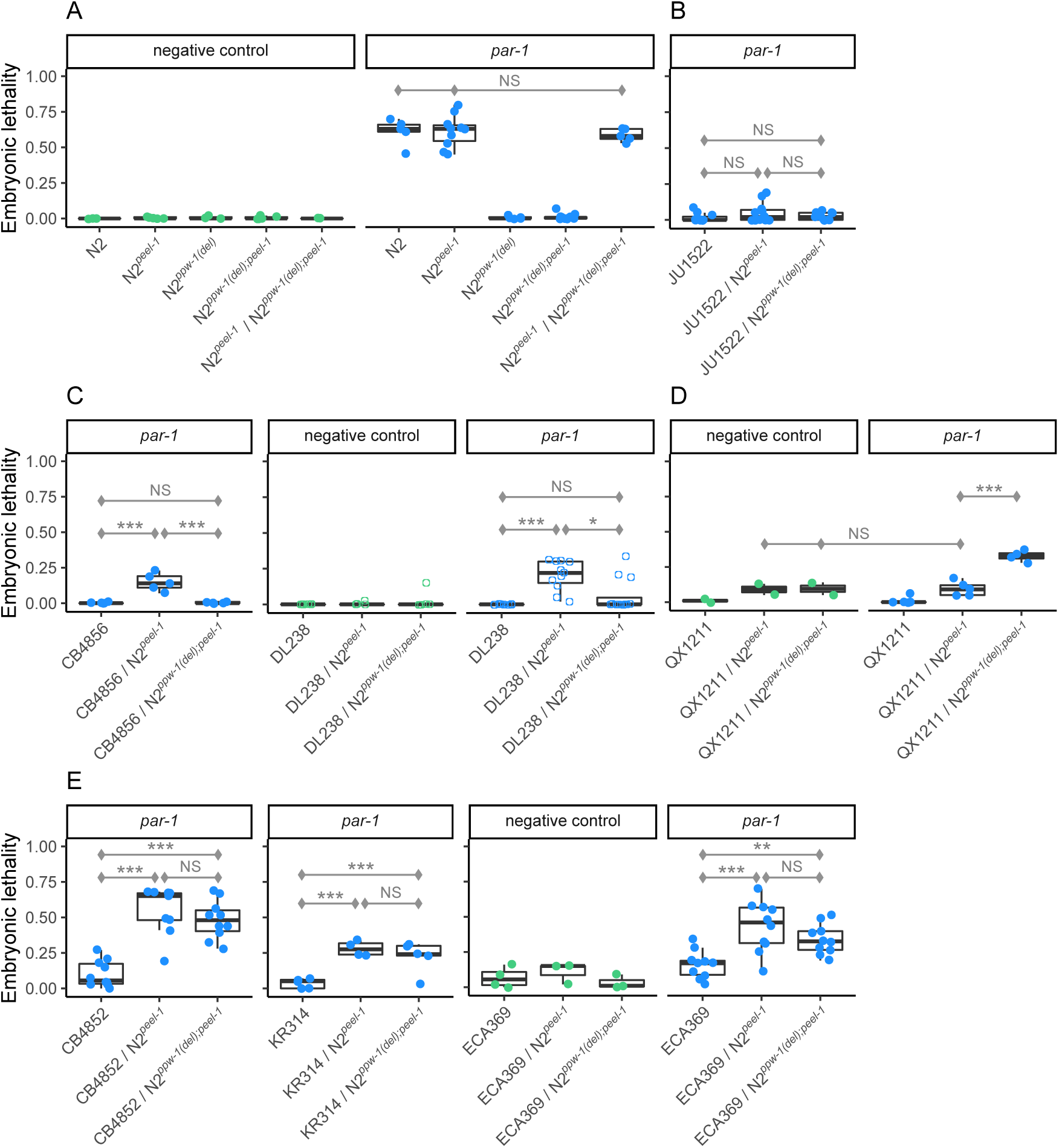
Complementation tests between seven wild isolates with low RNAi response and the RNAi-sensitive laboratory strain N2, with and without the *ppw-1* deletion allele. Response was measured by embryonic lethality following RNAi by feeding against the embryonic target *par-1*. Points represent the average across pooled hermaphrodites, with one exception (see below). (A) To circumvent embryonic lethality arising from the *zeel-1;peel-1* genetic incompatibility (*Seidel et al., 2008,* 2011), we used a null allele of the sperm-delivered toxin *peel-1* in N2, which has no effect on RNAi in either the responsive (N2) or the resistant (N2*^ppw-1(del)^*) backgrounds. A single copy of *ppw-1* is sufficient to fully restore the germline RNAi response in N2. (B) - (E) Complementation tests for seven wild isolates with weak germline RNAi, representing a diversity of genetic backgrounds. Embryonic lethality in the control condition is shown when it was observed to be greater than zero. For DL238 (C), the open circles represent the proportion of dead embryos per individual; the overall pattern qualitatively replicates that which we observed in pooled hermaphrodites (Figure S2; individuals shown here to highlight variability). For QX1211 (D), the *sup-35;pha-1* incompatibility (*Ben-David et al., 2017*) induced embryonic lethality, visible in the control condition and the genotype without the *ppw-1* deletion in the *par-1* treatment (see File S1 for details). Significance levels (Tukey’s contrasts): p<0.001 (***), p<0.01 (**), p<0.05 (*).

The seven strains exhibited four distinct response patterns: (i) no rescue, (ii) *ppw-1*-dependent rescue, (iii) *ppw-1*-dependent suppression, and (iv) *ppw-1*-independent rescue, described in detail below. These results indicate that within *C. elegans*, PPW-1 activity varies, PPW-1 activity differentially affects germline RNAi due to interaction with other varying factors, or both. The results further suggest that weak germline RNAi is multigenic within each strain, since rescued responses were all lower than N2 levels, indicating the presence of factors other than *ppw-1*.

i. Strain JU1522 showed no rescue, i.e., no improved RNAi response in either F1 genotype (Figure 3B). This suggests that weak RNAi in JU1522 is independent of *ppw-1*, or at least that alleles that promote RNAi in N2, including *ppw-1*, are not haplo-sufficient to increase the response in the JU1522 background.
ii. Strains CB4856 and DL238 exhibited *ppw-1*-dependent rescue: an increased RNAi response when crossed to N2, but only in the background with the wild-type *ppw-1* allele (Figure 3C, Figure S2). This outcome in CB4856 is consistent with prior reports (*Pollard & Rockman, 2013; Tijsterman et al., 2002*). In DL238, replicate to replicate variation in embryonic lethality was high, so we investigated whether this could be explained by potentially stochastic induction of the RNAi response between individual worms. This appears to be the case: tested individually, some hermaphrodites produced no dead embryos and others over 30% (Figure 3C).
iii. Unexpectedly, QX1211 showed *ppw-1*-dependent suppression: the heterozygote with the *ppw-1* deletion allele exhibited a significant increase in embryonic lethality, implying that reduction of *ppw-1* in this strain promotes germline RNAi (Figure 3D). As expected in this cross, we also observed lethality arising from the genetic incompatibility at the *sup-35;pha-1* locus (*Ben-David et al., 2017*) (details in File S2).
iv. Strains CB4852, KR314, and ECA369 exhibited *ppw-1*-independent rescue, in which the two heterozygote genotypes exhibited levels of embryonic lethality that were equivalent to each other and significantly higher than the wild isolate on its own. This suggests that N2 alleles other than *ppw-1* promote the RNAi response in these genetic backgrounds (Figure 3E).

To ensure that differences in lethality came from variation in RNAi genes and not from developmental variation specific to *par-1* (*Paaby et al., 2015*), we introgressed a germline-expressed GFP construct into four strains representing the four observed response patterns and quantified fluorescence following RNAi against GFP. With the exception of ECA369, which showed higher than expected RNAi sensitivity, the responses confirmed RNAi incompetency (Figure S3).

The results of these complementation tests demonstrate diversity in the function or effect of PPW-1 activity within *C. elegans*. Further, the *ppw-1*-independent rescue and the incompleteness of the *ppw-1*-dependent rescue implicate functional variation at genes other than (or in addition to) *ppw-1*, indicating that wild-type strains are likely to carry mutations affecting RNAi at multiple genes. However, with the exception of the unique *ppw-1* frameshift in CB4856, the extent to which RNAi alleles are likely to be strain-specific, versus shared across the population, is unclear.

### Genetic complementation between wild isolates implicates diverse and polygenic basis for germline RNAi incompetence

To examine whether alleles limiting germline RNAi are shared across strains, we crossed low-response strains to each other and looked for complementation. As above, we measured embryonic lethality following *par-1* RNAi exposure in the F1 generation. We performed a total of six tests, with each pairwise cross for strains CB4856, DL238, KR314 and QX1211 (Figure 4), chosen based on their diversity in PPW-1 function (Figure 3) and compatibility at the *zeel-1*;*peel-1* locus (*Andersen et al., 2012*) (see File S2 for more details).

**Figure 4.**
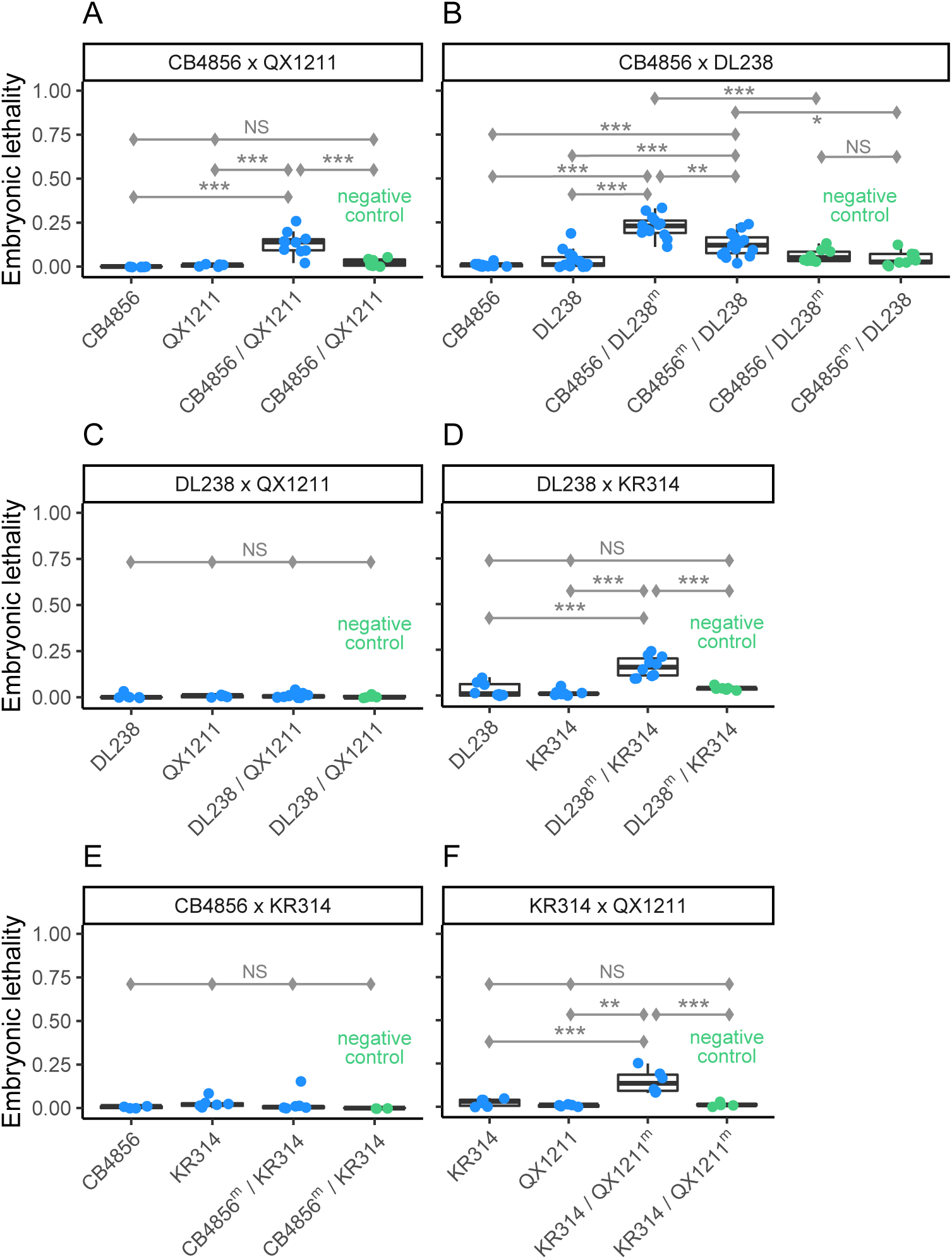
Pairwise complementation tests between four wild isolates with low RNAi response. (A) - (F) Response was measured by embryonic lethality following RNAi by feeding against the embryonic target *par-1*. The heterozygote genotypes were generated by crossing males and hermaphrodites in both directions (A) - (C), except for crosses with strain KR314, which does not produce fertile males (D) - (F). With the exception of CB4856 × DL238 (B), in which hermaphrodites sired by DL238 exhibited a significantly stronger response than those sired by CB4856, cross direction had no effect on embryonic lethality in the next generation and plots show pooled data. Significance levels (Tukey’s contrasts): p<0.001 (***), p<0.01 (**), p<0.05 (*).

We observed multiple instances of complementation, indicating that variation in RNAi is polygenic and that low-response strains carry alleles with distinct functional effects. For example, in the CB4856 × QX1211 cross, the heterozygote produced significantly more dead embryos than either strain on its own (Figure 4A), indicating that alleles that dampen the RNAi response are not shared since their function is at least partially rescued by the alternate genetic background. Given that CB4856 and QX1211 exhibit responses that are dependent on *ppw-1* but opposite to each other (Figure 3), their complementation may be occurring at *ppw-1* itself. However, despite similar responses under *ppw-1* manipulation (Figure 3), DL238 and CB4856 also complement (Figure 4B), suggesting distinct mechanisms. DL238 failed to complement QX1211 (Figure 4C), but did complement KR314 (Figure 4D), indicating shared and distinct mechanisms, respectively; these outcomes are the opposite of those observed for CB4856 crossed to the same strains (Figure 4A, 4E), reinforcing the conclusion that CB4856 and DL238 harbor distinct genetic mechanisms. We also saw evidence for distinct mechanisms in the complementation of KR314 × QX1211 (Figure 4F).

The rescued responses of the crossed strains point to pervasive diversity in the genetic mechanisms that underlie germline RNAi response in *C. elegans*. Together, the two sets of complementation assays (Figure 3, Figure 4) demonstrate that RNAi incompetence is multigenic within individual strains and caused by diverse alleles with distinct functional effects. This in turn indicates that variation in RNAi is a polygenic phenomenon within *C. elegans*, and suggests that it may be mediated by rare variants. In addition to *ppw-1*, causal mutations may reside in other Argonautes: though *ppw-1* is essential for germline RNAi in N2, overexpression of other WAGOs can rescue the response, implicating interchangeability (*Yigit et al., 2006*). We hypothesize that natural variation in the expression or function of RNAi genes, specifically WAGOs, produces a phenomenon of gene regulation by small RNAs that is highly diversified within the species. Variability in multiple factors is consistent with the dramatic range in sensitivity to germline RNAi overall (*Paaby et al., 2015*) as well as the diversity in genetic mechanisms underlying incompetence observed here.

### *C. elegans* RNAi genes show unusually high variation in expression

To assess whether and how RNAi genes might vary in expression across strains, we performed RNA-seq on low-response strains CB4856 and QX1211, on responsive strains N2 and JU1088, and on strain EG4348, which shows an intermediate response (*Paaby et al., 2015*). RNA samples were prepared from young, reproductively mature hermaphrodites without RNAi induction. Relative to other genes in the genome, RNAi genes, in particular Argonautes, showed highly elevated expression variation across strains, including *ppw-1* and the related WAGO *sago-2*.

First, we examined expression at 62 genes known to directly mediate RNAi (listed in Table 1), including Argonautes currently classified as pseudogenes on Wormbase (*Harris et al., 2020*). Of these, all but two were expressed in every strain: *wago-5* was not expressed at all, and the putative pseudogene *ZK218.8* was not expressed in the responsive stains but was expressed in CB4856 and QX1211. Of the genes with detectable transcripts, approximately half (29/61) exhibited differential expression (FDR<0.1) between N2 and at least one other wild strain (Figure 5A).

**Figure 5.**
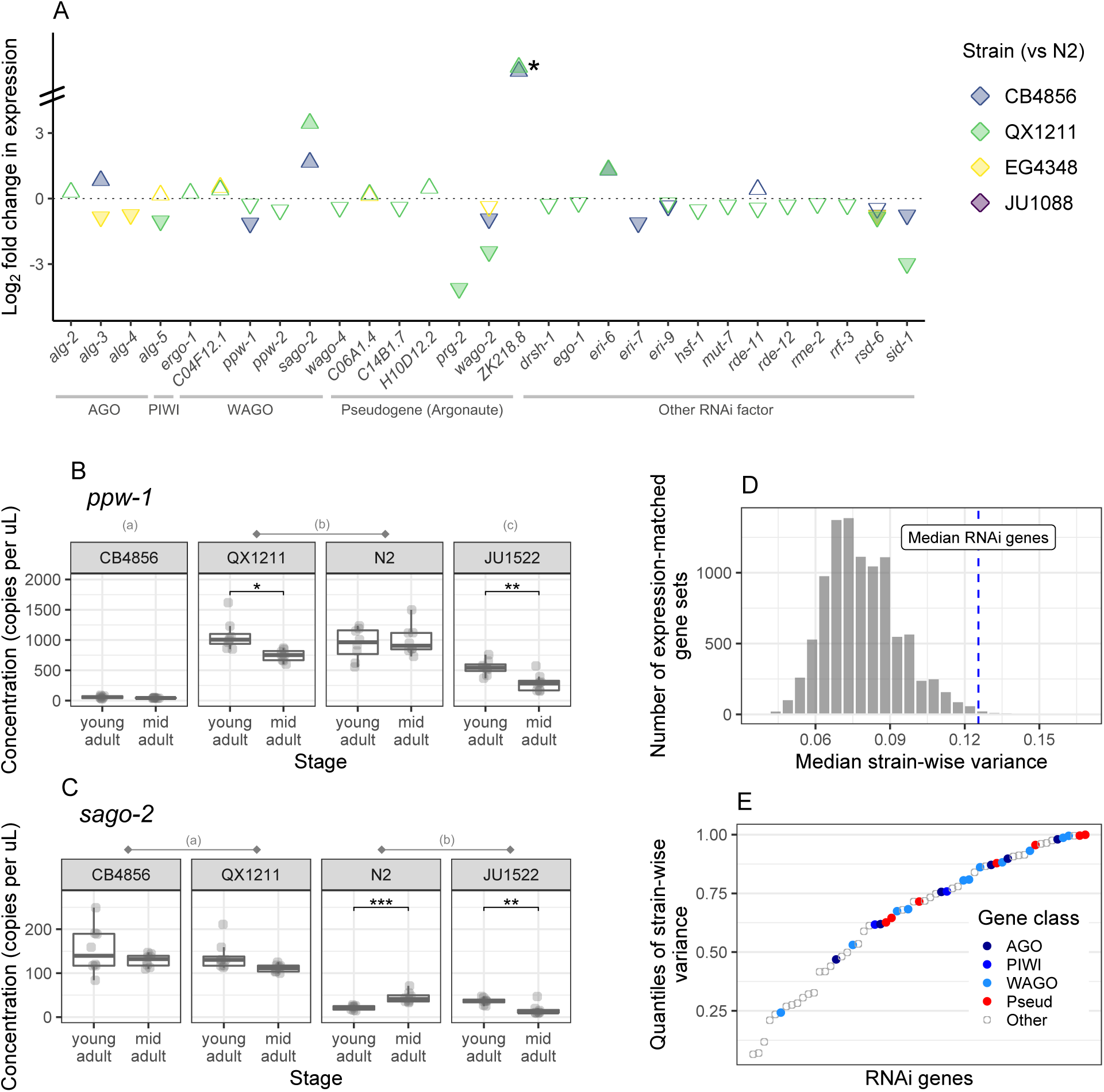
Variation in gene expression for RNAi factors. (A) Via RNA-seq, we examined 62 genes for differential expression between N2 and low-response strains CB4856 and QX1211, high-response strain JU1088, and moderately responsive strain EG4348. Only genes with significant results (FDR<0.1) are displayed; filled arrows indicate fold change >1.5. The (*) at *ZK218.8* indicates differential expression beyond the y-axis scale; this gene is not expressed in the N2 reference strain and has been classified as a pseudogene. (B) - (C) Expression differences via droplet digital PCR for *ppw-1* and *sago-2*. Ten strains were evaluated (Figure S4); a subset are shown here. Across strains, significant differences (Tukey’s contrasts, p<0.05) are indicated by letter groupings; for example, QX1211and N2 have equivalent concentrations of *ppw-1*, while CB4856 and JU1522 concentrations are significantly different from all others. Within each strain, significant differences between developmental stages (pairwise contrasts with Bonferroni correction) are indicated by: p<0.001 (***), p<0.01 (**), p<0.05 (*). (D) Histogram of median strain-wise variance for 10,000 gene sets, expression-matched to the RNAi genes, randomly sampled across the genome following variance-stabilizing transformation. (E) Strain-wise variance of the 61 expressed RNAi genes, plotted by quantile of genome-wide variances (gene expression data as in D). Argonautes, including those currently classified as pseudogenes, are indicated by filled circles.

**Table 1.**
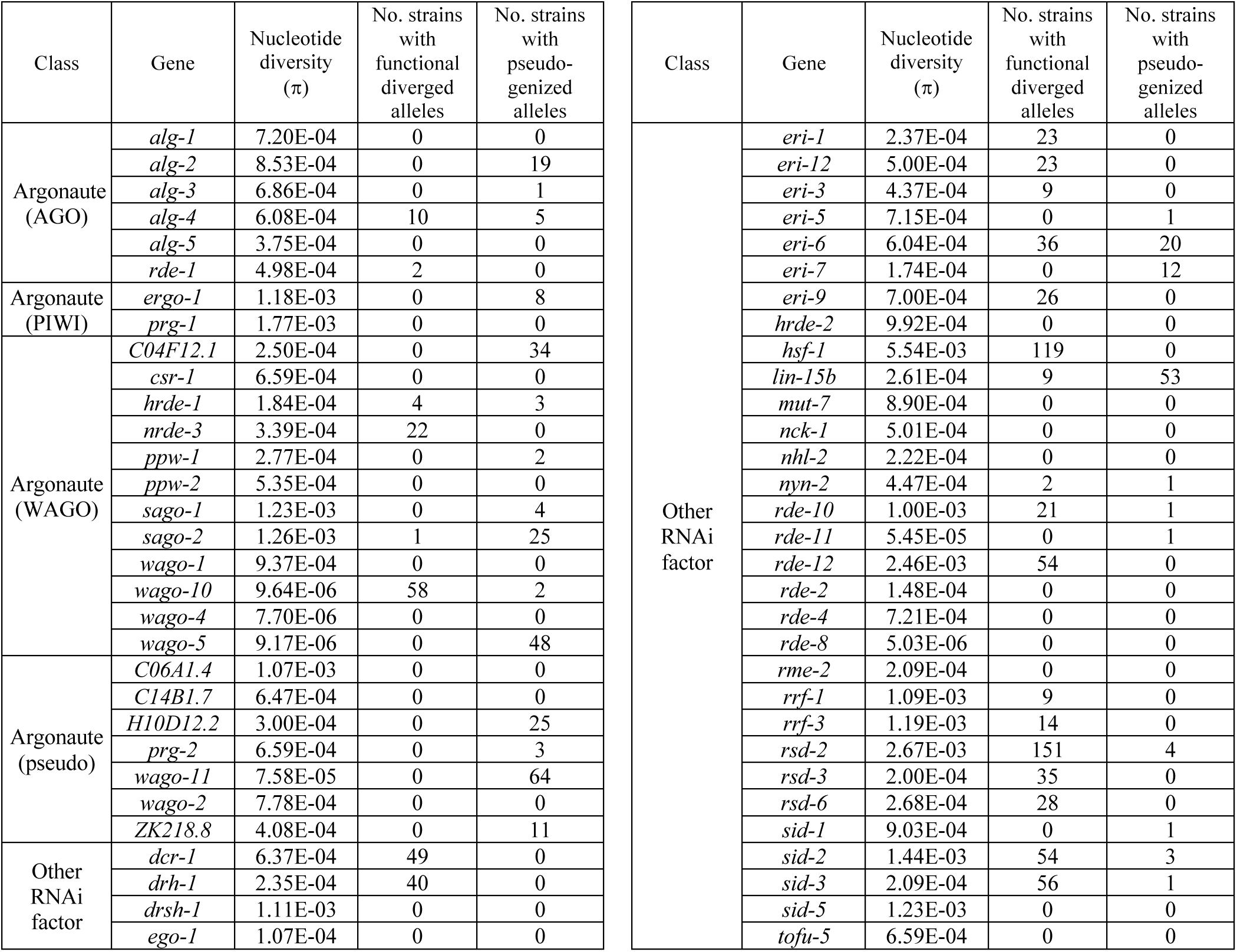
Molecular diversity at 62 RNAi genes, from 403 strain isotypes. Genes were manually curated from the literature and include 20 putatively functional Argonautes and seven currently classified as pseudogenes on Wormbase (*Harris et al., 2020*), though pseudogene status varies in the literature (and likely across strains). “Functional diverged alleles” are those with at least 1% nucleotide divergence from the reference genome, including at least five moderate mutations, such as amino acid substitutions, and no disruptive high impact mutations, such as frameshifts or stop-gains. “Pseudogenized alleles” are those with at least one high impact mutation called with high confidence and at which 1% or more of the sites are diverged or missing, or those with at least 50% of the sites with missing calls. Potential pseudogenization was assessed with respect to the reference genome and considered even for those Argonautes already classified as putative pseudogenes.

The strains with greatest differential expression were those with weakest germline RNAi (Figure 5A). That is: QX1211, then CB4856, showed the most differences across the gene set (26/61 and 10/61); the moderately responsive strain EG4348 showed a handful of differences (7/61); and the highly responsive strain JU1088 showed a difference at only one gene (*rsd-6*) (Figure 5A). Of genes differentially expressed by both CB4856 and QX1211, the direction of expression was concordant with one exception (*rde-11*), including reduced expression of *ppw-1* and elevated expression of *sago-2*. It is critical to note that though QX1211 and CB4856 show the greatest degree of differential expression relative to N2, they are also the most genetically diverged (*Cook et al., 2017*). That said, RNAi incompetence does not appear to be a function of genetic distance from the reference strain, as highly diverged isolates ECA701, JU561, and XZ1516 (*Crombie et al., 2019*) were responsive to *par-1* RNAi (data not shown), and RNAi sensitivity for 55 wild isolates in (*Paaby et al., 2015*) showed no relationship with divergence from N2 (Figure S4).

The WAGO *sago-2* shares high sequence identity with *ppw-1* and resides ∼17cM away on chromosome I. These two genes share overlapping function in the N2 background (*Yigit et al., 2006*), so the underexpression of *ppw-1* and overexpression of *sago-2* in CB4856 and QX1211 (Figure 5A) caught our attention. However, in some strains, including QX1211, poor mapping of short reads to the reference genome (*Cook et al., 2017*) at these loci suggests gene divergence or duplication. We resolved sequence ambiguities via de novo assembly of paired-end reads and long read sequencing, and observed that QX1211 carries *ppw-1*-like alleles at both loci (File S3). Therefore, to confirm our RNA-seq observations of *ppw-1* and *sago-2* expression and also to evaluate additional strains, we designed a droplet digital PCR (ddPCR) experiment to measure both transcripts simultaneously and discriminate between them using transcript-specific labels. We tested all strains thus far discussed: the seven low-response strains for which we tested *ppw-1* function (Figure 3), as well as responsive strains N2, JU1088, and EG4348. To evaluate whether *ppw-1* or *sago-2* expression changed as worms aged, we also assayed two developmental timepoints, young adult and mid adult.

The ddPCR results were consistent with our RNA-seq observations, and in sum confirm high variability in *ppw-1* and *sago-2* expression (Figure 5B-C, Figure S5). Overall, *ppw-1* expression was about an order of magnitude greater than that of *sago-2*, and both expression levels and changes in expression between developmental timepoints differed significantly across strains for both genes (Table S2). Taken individually, neither *ppw-1* nor *sago-2* expression correlated with RNAi responsiveness, and across all ten strains, the combined expression was both highest and lowest in two strongly resistant strains: QX1211 and JU1522, respectively (Figure 5B-C, Figure S5). JU1522 has consistently exhibited negligible germline RNAi, including no rescue when crossed with N2 (Figure 3) and no response even at later age (Figure S3); this strain shows the weakest response we have observed. One possibility is that both high and low expression of *ppw-1* and *sago-2* limit germline RNAi. This hypothesis fits with our observation that a haploid dose of *ppw-1* increases the RNAi response in QX1211 (Figure 3D), and with the prior finding that *ppw-1* and *sago-2* encode functionally interchangeable proteins that can compensate each other (*Yigit et al., 2006*). However, it is inconsistent with the observation that overexpression of these factors increases RNAi sensitivity in N2 (*Yigit et al., 2006*). N2 and JU1088, the two strains with the most robust germline RNAi response in our analysis, exhibited intermediate levels of *ppw-1* and *sago-2* combined (Figure S5).

Having observed significant expression variation for many of the RNAi genes, we next asked whether this gene set is more variable than other genes in the genome. The answer is yes: for the 61 actively transcribed RNAi genes, the median strain-wise variance (after variance-stabilizing transformation, see Methods) was higher than that of expression-matched sets randomly sampled from the genome 99% of the time (9936/10,000 comparisons, Figure 5D); 74% of these (7351/10,000) were statistically significant (one-tailed Mann-Whitney test, alpha=0.05), far exceeding that expected by chance. This effect is driven by elevated variance across the gene set, not by a few outliers of high variance, as the majority (57%) (35/61) exhibit significant differences by strain via liklihood ratio test (FDR<0.1) compared to 29% of all genes in the genome. Thus, RNAi genes are highly enriched for strain-wise variation (p=4.8x10^-6^, hypergeometric test).

Several of the RNAi genes are exceptionally variable. The putatively pseudogenized Argonaute *prg-2* exhibits the 10^th^ highest strain-wise variance in the genome, and six genes, including *ppw-1* and *sago-2*, are in the top 2%. Given the relatively recent evolution of WAGOs (*Buck & Blaxter, 2013*) and their potential redundancy in function (*Yigit et al., 2006*), we expected to see especially high expression variation for this gene class. This hypothesis is well supported: all but one (*sago-1*) of the 12 WAGOs are in the top half of genome-wide variance, though Argonautes of all classes showed a similar trend, including those presumed to be pseudogenes (Figure 5E). The distribution of strain-wise variances was more evenly distributed for other RNAi factors and included both highly variable and highly invariant expression patterns (Figure 5E).

The elevated expression variation in RNAi genes represents heritable variation in small RNA processes in *C. elegans*, consistent with the other evidence for diversification of germline RNAi function within the species and a possible explanation for that diversity. Variable expression of putative pseudogenes, including the active transcription of *ZK218.8* in some non-reference strains, also prompts the question of whether wild-type strains vary in their complement of functional Argonautes. *ZK218.8* was previously identified as an Argonaute (*Yigit, 2007*) but remains unexplored in the literature; its expression signature suggests that pseudogenization may have occurred in some strain lineages but not in others. While these observations all point to diversification in genetic mechanisms of RNAi, the historical forces driving these outcomes remain obscured. To evaluate this, and to look for evidence of strain-specific mutations affecting RNAi, we next turned to population-level sequence data.

### *C. elegans* RNAi genes show lineage-specific diversification and pseudogenization

We examined allelic diversity at the 62 RNAi genes, both to identify candidate mutations for RNAi incompetence and to assess selection history via patterns of molecular variation. We examined genotypes in CeNDR, which includes hundreds of strains representing the global *C. elegans* population (*Cook et al., 2017*). For the seven low-response strains tested in the lab, we identified putatively deleterious variants that may contribute to individualized loss of function, as many were strain-specific (Table S3). Across the population, we observed substantial sequence variation among all gene classes, including strain-specific instances of pseudogenization and allelic divergence (Table 1). These results indicate that many RNAi genes have recently undergone rapid and dynamic evolution, by what appears to be both relaxed and intense selection pressure. The occurrence of rare alleles and lineage-specific patterns support the functional diversification we observe in the lab and support a model of dynamic contemporaneous evolution of small RNA pathways in *C. elegans*.

Relative to genome-wide averages (*Lee et al., 2021*), estimates of nucleotide diversity (π) per gene were generally high (Table 1), with π>0.001 for 14/62 genes (13/55 genes not characterized as pseudogenes). Elevated variation was often, but not always, associated with high-impact mutations in one or more strains (Figure S6). Therefore, to distinguish between instances of pseudogenization and gene loss versus functional allelic diversification arising from directional or balancing selection, we classified highly diverged alleles two ways. We considered an allele to be pseudogenized if it harbored at least one high impact mutation called with high confidence and if at least 1% of the sites were diverged or missing relative to the reference genome, or if over 50% of sites were missed calls; we classified “functional diverged alleles” as those with at least five amino acid substitutions, at least 1% divergence across sites, and no high impact mutations called at the locus.

We observed extensive pseudogenization across the gene set. Given the expansion and diversification of Argonautes in nematodes (*Buck & Blaxter, 2013*), we hypothesized that the WAGOs might be relatively unconstrained and therefore especially susceptible to gene loss. Indeed, many WAGOs (7/13) showed evidence of pseudogenization, but putative loss occurred in all gene classes, including the PIWIs, AGOs, and other RNAi factors (Table 1). For example, the PIWI Argonaute *ergo-1* exhibits extensive variation and lineage-specific pseudogenization, indicative of relaxed selection (Figure 6A). Excluding the seven Argonautes classified as pseudogenes on Wormbase (*Harris et al., 2020*), 40% (22/55) of genes indicated pseudogenization in one or more strains (Table 1). In contrast, a few genes exhibited very low polymorphism (Figure 6B, Figure S6), likely reflecting evolutionary constraint and purifying selection. These genes were mostly non-Argonautes, with the exception of *csr-1* and *prg-1*, the only Argonautes essential for development (*Yigit et al., 2006*).

**Figure 6.**
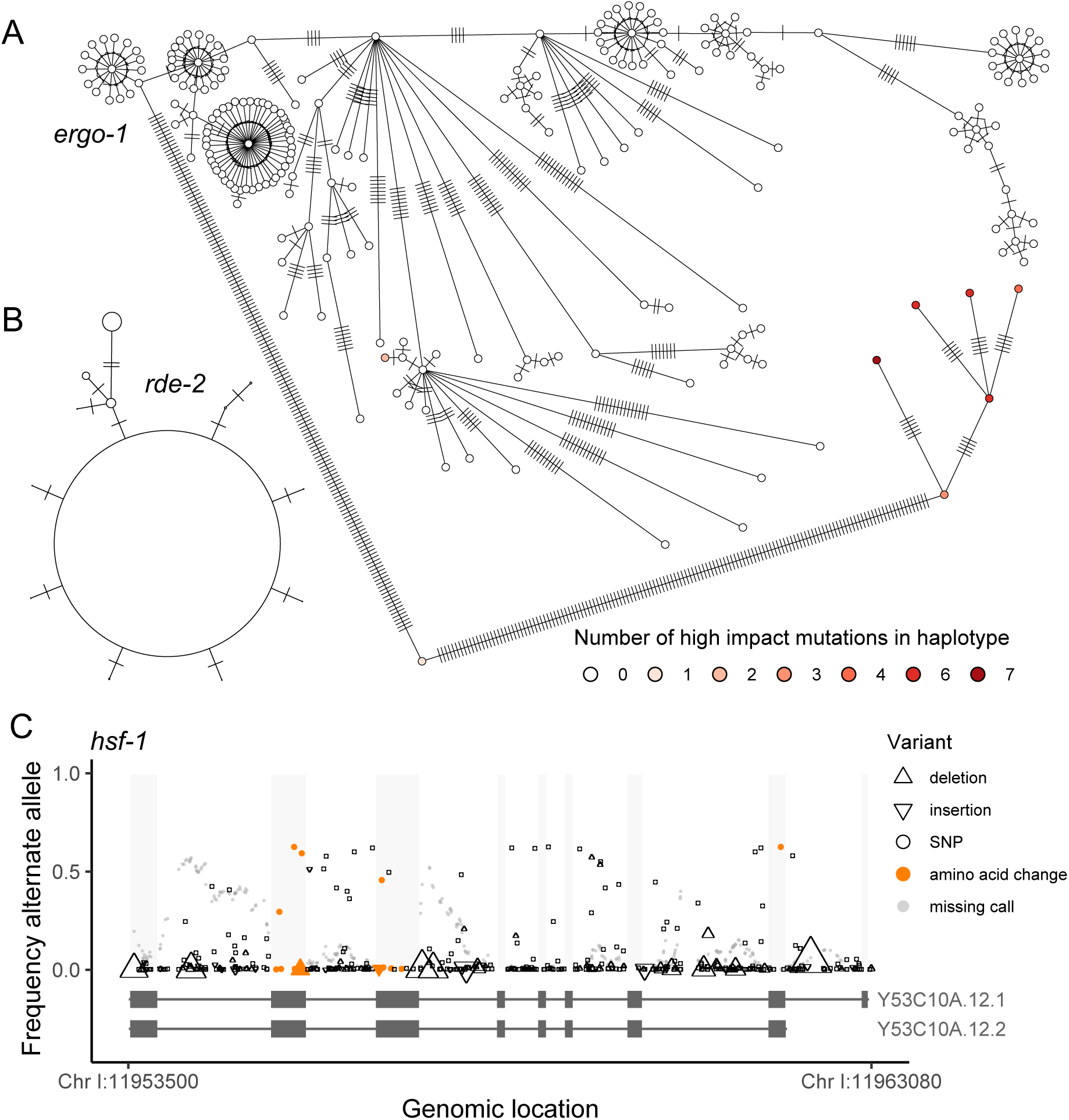
Population-level allelic diversity in RNAi genes across 403 strains. (A) The Argonaute *ergo-1* exhibits elevated polymorphism. Accumulation of high impact mutations in some alleles indicates gene loss by pseudogenization and implies relaxed selection. In this haplotype network, each circle represents a unique haplotype (of one or more strains) and hatch marks indicate mutations. (B) In contrast, *rde-2* exhibits conservation. Most strains belong to the common haplotype; haplotype circles are scaled by frequency. (C) The transcription factor *hsf-1* also shows elevated polymorphism. Multiple changes to the amino acid sequence, mutations segregating at intermediate frequency, and no observed high impact mutations that disrupt the protein suggest functional divergence. Each observed mutation is represented as a single point; the up- and down-triangles representing indels are scaled by indel length (range = 1-99, mean = 6.8).

We also observed pervasive functional divergence, with functionally diverged alleles in 25/62 genes (Table 1). Unlike the pseudogenized alleles, which often occurred singly, the functional diverged alleles were most often shared across strains, consistent with positive selection driving or maintaining divergent gene function. For example, *hsf-1* exhibited very high polymorphism, including amino acid changes at intermediate frequencies, but no instances of high impact mutations likely to knock out function (Figure 6C). (We also observed poor read mapping across the locus, which indicates further divergence but may also obscure deleterious mutations.) This transcription factor is a master regulator of other RNAi genes, and HSF-1 activity is associated with transgenerational inheritance of an on/off RNAi response (*Houri-Zeevi et al., 2020*). As *hsf-1* is a potential keystone regulator of small RNA pathways, its diversification may underlie significant functional variation in RNAi.

The patterns of elevated polymorphism indicate that RNAi genes are evolving dynamically within *C. elegans*, with lineage-specific trajectories of relaxed selection and gene loss, as well as possible directional selection and functional divergence. These findings are consistent with our experimental observations that the genetic basis of RNAi failure is strain-specific. Moreover, they suggest that functional characterizations of these genes, which have been universally achieved in N2, may be strain-specific as well.

## DISCUSSION

In this study, we demonstrate that a diversity of genetic mechanisms underpins the failure of some wild-type *C. elegans* strains to mount a robust germline RNAi response. Rather than identifying one or more common, shared factors that explain RNAi incompetence, our results indicate that RNAi fails for different reasons in different strains and that the same genes can produce opposite responses. Coupled with high levels of divergence and lineage-specific pseudogenization at known RNAi genes, these findings indicate that the small RNA pathways in *C. elegans* are evolving rapidly and dynamically, leading to functional diversification of RNAi activity.

We propose that such diversification evolved as a consequence of (a) redundancy and interchangeability among Argonautes (*Billi et al., 2014; Yigit et al., 2006*), (b) competition between overlapping pathways (*Yigit et al., 2006; Youngman & Claycomb, 2014*), and (c) a population structure with reduced gene flow (*Dolgin et al., 2007*). Small RNA processes dominate the biology of *C. elegans* (*Houri-Zeevi et al., 2020; Youngman & Claycomb, 2014*), and defenses against pathogens and transposable elements may be especially susceptible to strong selection (*Nuez & Félix, 2012*). However, because the species is globally dispersed and reproduces primarily by selfing, *C. elegans* lineages evolve semi-independently, may be exposed to distinct selection pressures, and may accumulate co-adapted allelic combinations (*Campbell et al., 2018; Dolgin et al., 2007*). Argonautes and other factors are shared among pathways, and competition between exogenous and endogenous RNAi can force induction of one pathway over another (*Yigit et al., 2006; Youngman & Claycomb, 2014*). In this vein, we might imagine, for example, how selection on germline maintenance in one genetic background could compromise a response to environmental triggers, as well as how Argonaute redundancy could facilitate evolutionary lability and gene-specific, lineage-specific responses even under similar selection pressures.

An example of competition between overlapping pathways may be reflected in some of our results, as simultaneous exogenous and endogenous demands on shared factors may explain the behavior of QX1211. In addition to the on/off responses among individuals (Figure 1, Figure 2), QX1211 exhibits increased sensitivity to germline RNAi after consecutive generations at ≥18°C (unpublished data)—but this co-occurs with onset of the mortal germline phenotype and reproductive extinction, which is associated with shifts in piRNA-like pools of small RNAs (*Frézal et al., 2018*). Thus, changes in RNAi activity appear to be either a cause or a consequence of germline mortality in QX1211 within individual animals, which in turn may explain the differences in sensitivity to exogenous RNAi. Competition between pathways may also explain the *ppw-1*-dependent suppression of RNAi in QX1211 (Figure 3). If *ppw-1* is a limiting factor in the defense against germline mortality in QX1211, then decreasing its availability might downregulate germline protection while simultaneously releasing resources for the competing exogenous pathway. QX1211 exhibited highest expression of *ppw-1* and *sago-2* (Figure 5) and carries a *ppw-1*-like allele of *sago-2* (File S3)—in the most speculative case, this might reflect a history of selection for *increased* germline-associated RNAi response in QX1211, even as laboratory assays for exogenous RNAi reveal apparent incompetence.

Consequently, although incompetence for laboratory-induced RNAi is the explicit focus of this study, we emphasize that the synthetic phenomenon of RNAi by feeding does not necessarily represent processes most relevant in nature. The role of RNAi in the wild remains largely obscured, though some observations offer clues. Exogenous RNAi likely induces responses that evolved for antiviral immunity, as viruses that infect *C. elegans* and other *Caenorhabditis* species have been discovered, notably in isolates with defective RNAi, and antiviral immunity shows a clear association with an active RNAi response (*Félix et al., 2011; Sarkies et al., 2013; Schott et al., 2005; Wilkins et al., 2005; Yigit et al., 2006*). The overlap between experimental RNAi and antiviral response is incomplete, however, as variation in RNAi sensitivity does not completely correlate with immunity and the systemic and transgenerational properties of RNAi are not observed in viral infection (*Ashe et al., 2013,* 2015*; Félix et al., 2011*). Orsay virus, the only naturally-occurring virus known to infect *C. elegans*, invades intestinal cells and is horizontally, but not vertically, transmitted (*Félix et al., 2011; Franz et al., 2014*), though vertically transmissible viral-like RNAs have been detected in the germlines of wild-caught *Caenorhabditis* isolates (*Richaud et al., 2019*), suggesting undiscovered host-pathogen dynamics. Endogenous RNAi is likely required for germline maintenance in the wild, as suggested by the observations in QX1211. Hence the piRNA pathway, which is active in the germline and presumed critical for maintaining genome integrity (*Wilson & Doudna, 2013; Youngman & Claycomb, 2014*), may dominate the biology of, or be upregulated more often in, some strains relative to others (*Frézal et al., 2018*). One possibility is that *ergo-1* gatekeeps RNAi pathway activity differently in different isolates, as *ergo-1* N2 mutants show enhanced exo-RNAi but reduced endo-RNAi (*Yigit et al., 2006*) and the *ergo-1* locus exhibits extreme allelic diversification in nature (Table 1, Figure 6). Another possibility is that in the wild, RNAi in the germline matters most to future generations. Strains resistant to RNAi upon exposure can show transgenerational sensitivity (*Tijsterman et al., 2002*), and a growing body of research emphasizes the outsized role of RNAi in transgenerational inheritance (*Houri-Zeevi et al., 2020,* 2021). Thus, RNAi as we have studied it in the lab provides an oblique view into its role in nature.

Given the essentiality of RNAi to numerous biological processes—and the centrality of *C. elegans* in RNAi research—is the variation in this system surprising? Perhaps not: even as RNAi genes are shared deeply within the eukaryotic lineage (*Shabalina & Koonin, 2008; Wynant et al., 2017*), the variation we describe here mirrors patterns of RNAi incompetence and molecular evolution over longer timescales. The ability to silence genes by dsRNA appears intermittently, and shows evidence of rapid evolution within the *Caenorhabditis* genus (*Nuez & Félix, 2012; Winston et al., 2007*), across nematodes generally (*Buck & Blaxter, 2013; Dalzell et al., 2011*), and in other systems (*Obbard et al., 2009*). Argonautes and associated RNAi factors also exhibit taxon-specific patterns of gene duplication, loss, and diversification, likely representative of diversification of biological functions (*Buck & Blaxter, 2013; Dalzell et al., 2011; Obbard et al., 2009*). In other words, the contemporaneous variation in RNAi observed within extant *C. elegans* mirrors the great lability in RNAi observed over long timescales (*Nuez & Félix, 2012*).

*C. elegans* increasingly appears to be dominated by transgenerationally inherited small RNA programs (*Houri-Zeevi et al., 2020,* 2021) that vary significantly in nature (*Frézal et al., 2018*). This variation offers leverage: characterization of wild-type variation can elucidate condition-dependent mechanisms (*Chandler et al., 2013*), which are rampant in RNAi, a complex and intricate collection of interactions susceptible to unexpected outcomes (*De-Souza et al., 2019*) and sensitive to environmental conditions (*Houri-Zeevi et al., 2021*). Identifying mechanisms of variation will help to bridge the gulf between our understanding of the genetics of RNAi and the role of RNAi in nature, and future work may benefit from evaluating wild-type isolates in the context of carefully chosen environmental perturbations (*Rockman, 2008*). For example, temperature likely matters for RNAi, given the exquisite sensitivity of *C. elegans* to temperature (*Testa et al., 2020*) and the intimate relationship between temperature and other stresses and RNAi (*Frézal et al., 2018; Houri-Zeevi et al., 2021; Pagliuso et al., 2021*); our observations of QX1211 would have been obscured without rigorous temperature control. Sydney Brenner’s selection of *C. elegans* as a model species, and N2 as the strain of study, was fortuitous for the future discovery of RNAi (*Félix, 2008*), and it remains the most fertile area for elucidating gene regulation by small RNAs (*Youngman & Claycomb, 2014*). Now, characterizations of significant natural genetic and functional variation in RNAi provide a new access point for expanding our understanding in a system already so well established.

## MATERIALS AND METHODS

### Strains used in this study

Table S4 contains a complete list of strains used in this study. To introduce germline-expressed GFP into wild isolates, we introgressed *zuIs178 [his-72(1kb 5’ UTR)::his-72::SRPVAT::GFP::his-72 (1KB 3’ UTR) + 5.7 kb XbaI -HindIII unc-119(+)]; stIs10024 [pie-1::H2B::GFP::pie-1 3’ UTR + unc-119(+)]* into strains CB4856, ECA369, JU1522 and QX1211 by crossing to RW10029 and backcrossing to the wild strain for 10-18 generations.

### Worm husbandry

Worms were cultured following standard protocol (*Stiernagle, 2006*), though we added 1.25% agarose to plates used to maintain non-N2 wild isolates, to avoid burrowing. Worms were maintained at 20°C without starving for at least three generations before initiating an experiment, with the exception of QX1211, which was maintained at 18°C to avoid induction of the mortal germline phenotype (*Frézal et al., 2018*).

### RNA interference

#### General culture conditions

RNAi was induced by feeding and experiments were carried out on plates, at 20°C, based on methods previously described (*Ahringer, 2006; Kamath et al., 2001*). In brief: to target endogenous germline-expressed genes, we fed worms HT115 *E. coli* bacteria that had been transformed with the pL4440-derived *par-1* (H39E23.1), *par-4* (Y59A8B.14), *pos-1* (F52E1.1), or GFP feeding vector (*Timmons et al., 2001*). The *par-1* and *pos-1* vectors were obtained from the Ahringer feeding library (*Kamath & Ahringer, 2003*); *par-4* was a gift from M. Mana. To target GFP, we transformed HT115 with pL4417, which carries 0.7 kb of GFP coding sequence (*Timmons et al., 2001*). We used *E. coli* carrying the empty pL4440 vector as a negative control. Bacteria were streaked from frozen stocks onto LB agar plates with carbenicillin (25 ug/mL) and tetracycline (12.5 mg/mL); liquid cultures were inoculated with 5-10 colonies from <1 week old plates, into LB broth with carbenicillin (50 ug/mL) and tetracycline (12.5 mg/mL) and incubated for 16-18hrs shaking at 37°C, then amplified in a 1:200 dilution with carbenicillin (50 ug/mL) for 6hrs. Seeded plates were incubated in the dark at room temperature and used no earlier than 44hrs and no later than 78hrs. Experimental worms were exposed to RNAi bacteria as L1s by hatching on RNAi plates, synchronized either by bleaching (*Stiernagle, 2006*) or by timed egg-laying by the hermaphrodite mothers.

#### Embryonic lethality assays

To measure RNAi response by phenotypic penetrance, we targeted *par-1* or *pos-1* transcripts in the hermaphrodite germline and measured embryonic lethality in the next generation.

Experimental worms were reared on RNAi plates and transferred as L4s to fresh RNAi plates for the egg-laying assay, remaining continuously exposed to RNAi bacteria since their hatching. For all experiments except those explicitly testing variation in penetrance between individual worms (Figure 1B, Figure 3C), the L4 hermaphrodites were pooled in small groups of 4-6 on 6-10 replicate assay plates. For the complementation tests (Figure 3, Figure 4, Figure S2), all embryos within the first ∼15hrs of egg-laying were scored for hatching, typically 100-200 embryos per plate. For assays testing RNAi within a defined window of reproductive maturity, we scored the embryos laid in a 4-6hr window within the first 8hrs of egg-laying (Figure 1A), or a 2hr window 4hrs after egg-laying began (Figure 2C). For the experiment measuring RNAi in individual worms over their reproductive lifespan (Figure 1B, Figure S1), the L4 hermaphrodites were singled to RNAi plates, permitted to lay embryos, and continually transferred to fresh plates until they ceased to lay, or laid only unfertilized eggs. To score embryos as dead or alive, we removed the egg-laying adult(s), incubated the plates at 20°C for 24hrs, and counted (dead) embryos and hatched larvae using a stereoscope. Experiments included 6-10 (RNAi treatment) or 4-6 (negative control) replicate plates.

To test the effect of genotype on embryonic lethality following exposure to RNAi, the counts of dead embryos and hatched larvae from each replicate plate were bound together as a single response variable and modeled with a generalized linear model with a quasibinomial error structure, implemented by the *glm* function in R. The model included a single linear predictor for genotype and took the form E(Y)=g^−1^(β_0_+β_genotype_). Within each experiment, differences between specific genotypes were assessed by pairwise contrasts using the “Tukey” specification in the function *glht* in the R package *multcomp* (*Hothorn et al., 2008*).

#### RNAi against GFP

To measure germline RNAi by GFP knockdown, worms carrying a histone-linked GFP driven by a *pie-1* promoter were fed RNAi bacteria targeting GFP. Synchronized animals were grown on RNAi plates, then individually selected for imaging at the following stages: young adults (6±2hrs after exiting L4 stage), day one adults (24±2hrs), and day 2 adults (48±2hrs). For whole worm fluorescence imaging, animals were anesthetized with 10mM NaN_3_ and mounted on 2% agarose pads, then imaged using a 10× objective with the PerkinElmer UltraVIEW VoX spinning disk confocal microscope equipped with an EM-CCD camera. Raw images were exported as OME.TIFF files. We used Fiji (*Schindelin et al., 2012*) to acquire the sums of intensity in the Z projection, then quantitated the GFP fluorescence by subtracting the integrated intensity of the background, over the area of the worm, from the integrated intensity of the whole animal. To test whether RNAi-treated worms exhibited reduced fluorescence relative to control worms, we analyzed the six samples (three treatment timepoints, three control timepoints) for each strain using a one-way ANOVA, then performed treatment-control contrasts within each timepoint using the R function *TukeyHSD()*.

### Single-molecule fluorescence *in situ* hybridization

#### Sample preparation and imaging

Custom Stellaris FISH probes were designed with the Stellaris Probe Designer (LGC Biosearch Technologies). We excluded polymorphic sites during probe design. Worms were synchronized on tryptone-free NGM agar plates at the L1 stage and reared on RNAi bacteria as described above. Embryos were extracted by standard bleaching/washing, fixed using 3.7% formaldehyde in RNase-free phosphate buffered saline, and hybridized (100nM at 37°C for 4hrs) with a Quasar 570 labeled probe set targeting either *par-1* or *par-4*, following the manufacturer’s instructions. Samples were mounted using VECTASHIELD antifade mounting medium with DAPI (Vector Labs #H-1200) on no. 1 cover slides. Images were captured with a 100X oil immersion objective on a PerkinElmer UltraVIEW VoX spinning disk confocal microscope equipped with an EM-CCD camera and piezoelectric motorized stage. Three-dimensional image stacks were collected using Volocity 3D visualization software (PerkinElmer) and exported as TIFF files.

#### Quantitative Analysis

Image segmentation masks were applied and chromosome clusters were counted using ImageJ (*Schneider et al., 2012*). Quantification of single molecule FISH spots was performed using Aro, a MATLAB-based, machine learning pipeline designed for single-molecule visualization in worm embryos (*Wu & Rifkin, 2015*). The training sets for the random forest classifier were generated from multiple samples of each genetic background and treatment. To test whether means or variances in transcript counts differed for RNAi-treated versus untreated samples within a strain, we applied two-sample *t*-tests and *F*-tests, respectively. For these tests, we only considered early stage embryos (up to four cells). To evaluate changes in transcript abundance over a wider range of embryonic development, we considered embryos with up to 30 nuclei and used ANCOVA to ask whether, adjusted for embryo stage, transcript levels varied across strains within the negative control condition; and whether, adjusted for embryo stage, transcript levels varied between control and treatment conditions within a strain. We used minimal model selection to test for changes in the way transcript level depended upon embryo stage (i.e. changes in slope). We estimated ANCOVA effect sizes as ω^2^ using the R package *sjstats* (*Lüdecke, 2018*).

### RNA-seq

#### Library preparation and sequencing

Healthy cultures of strains N2, CB4856, QX1211, JU1088 and EG4348, reared for several generations without starving or bleaching, were bleached to retrieve large numbers of embryos. Synchronized L1 larvae were reared on plates with the empty RNAi feeding vector, details as described above. Young, reproductively mature hermaphrodites were washed off plates and rinsed twice with M9, then RNA was extracted with TRIzol (Invitrogen #15596026) and RNeasy columns (Qiagen #74104), following (*He, 2011*). All samples were collected and processed simultaneously and in triplicate, starting with replicate plates of worms. Libraries were prepared with the NEBNext Ultra II Directional RNA Library Prep Kit for Illumina (NEB #7760), with cDNA generated from fresh RNA samples using 500ng of RNA and 10 cycles of PCR. Libraries were quality checked using an Agilent 2100 Bioanalyzer and fragments were size-selected via BluePippon (Sage Science). Libraries were sequenced on an Illumina NextSeq for single-end 75bp reads at the Molecular Evolution Core facility at the Georgia Institute of Technology.

#### Alignment and gene expression quantification

We generated strain-specific transcriptomes for RNA-seq read quantification by patching SNPs and INDELs from CeNDR (release 20210121) (*Cook et al., 2017*) onto the N2 reference genome (release ws276) (*Harris et al., 2020*) using *g2gtools* (v0.1.31 via conda v4.7.12, Python v2.7.16) (https://github.com/churchill-lab/g2gtools), followed by transcriptome extraction. Specifically, for each non-reference strain, INDELs were first chained onto the reference genome using *g2gtools vcf2chain* and SNPs were patched onto the reference genome FASTA using *g2gtools patch*. Next, INDELs were chained onto the SNP-patched genome using *g2gtools transform* and strain-specific GTFs were created from this updated genome FASTA via *g2gtools convert*. Strain-specific transcriptomes were generated from these strain-specific genome FASTAs and GTFs using *gffread* (v0.12.7) (*Pertea & Pertea, 2020*).

Transcript-level quantification was performed using Salmon (v1.4.0) (*Patro et al., 2017*). Before Salmon quantification, Illumina TruSeq adapters were trimmed from RNA-seq reads using Trimmomatic (v0.3.9) (*Bolger et al., 2014*) with parameters *ILLUMINACLIP:TruSeq3-SE.fa:1:30:12*. Salmon index files were built from the strain-specific transcriptomes using command *salmon index* with options *-k 31 --keepDuplicates* (all others default; no decoy was used). Transcript quantification was performed with *salmon quant* with options *-l SR --dumpEq, --rangeFactorizationBins 4, --seqBias, and --gcBias*, and the library-specific fragment length arguments *--fldMean* and *--fldSD*.

#### Analysis of gene expression

We performed all expression analyses in R (v4.0.3) (*R Core Team, 2020*) using data processed with the DESeq2 package (v1.32.0) (*Love et al., 2014*). We used the *tximport* package (v1.20.0) (*Soneson et al., 2015*) to import Salmon transcript quantification data into DESeq2 and to compute gene-level expression quantification estimates. Genes with 10 or fewer counts total across all samples after *tximport* were excluded from downstream analysis (18,589 genes retained).

To test for differential expression across strains, gene counts were modeled using the negative binomial generalized linear model in DESeq2:

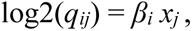

where for gene *i*, sample *j*, *q* is proportional to the true concentration of RNA fragments for the gene. *β_i_* gives the log2 fold changes for gene *i* and *x* represents the strain; batch was not included in the model because all samples (three biological replicates per strain, five strains) were processed simultaneously. Significance testing for differential expression was performed by likelihood ratio test (LRT) in DESeq2, which captured strain-wise significance by comparing the above model to a reduced model containing only the intercept (*Love et al., 2014*). Genes were considered differentially expressed by strain if the genome-wide adjusted p-value (FDR) from the LRT was <0.1 (5,464 of the 18,589 genes passed this threshold overall). Estimates of differential expression between N2 and each other strain were extracted via pairwise contrasts; effect sizes and p-values were corrected using the ‘*ashr’* method from the *ashr* package (v2.2-47) *(Stephens, 2017*).

We assessed strain-wise variance per gene by first obtaining normalized gene expression data from the variance-stabilizing transformation (*vst* function) in DESeq2. This transformation puts the data in log2 scale, incorporates library size and gene length normalizations, and makes the variance independent of the mean (*Love et al., 2014*). Strain-wise variance for each gene was then estimated by one-way ANOVA, i.e. *counts ∼ strain*; the sums of squares for the strain term was extracted using the *aov* function.

Expression-matched gene sets for the RNAi genes were constructed by first identifying, for each RNAi gene, all genes with *vst*-normalized mean expression (across all samples) within one percentile of the RNAi gene’s mean expression (+/- 0.005 in expression quartile). One of these genes was chosen at random for each RNAi gene, and this procedure was repeated 10,000 times to obtain the 10,000 random expression-matched gene sets. The median strain-wise variance (of *vst*-normalized gene counts) for each random set of genes was computed and compared to the median strain-wise variance of the RNAi genes.

### Droplet digital PCR

To design an unbiased primer set, we identified regions of identical sequence between *ppw-1* and *sago-2* and across the ten strains of interest. Following (*Kamitaki et al., 2018*), we chose primers to target both genes, and probes to discriminate between *ppw-1* (FAM) and *sago-2* (HEX). Sequences are as follows: forward (CTTGGTACCGCTCCGCTC), reverse (GCTGATTCGGTTTGATCGTC), *ppw-1* probe (AGACGAGAAATGTGGAGAGGGGAA), *sago-2* probe (AGACGAGAAATGAGGAGTGGGGAA). Both probes anneal in the same location, ensuring competition between them.

Worms from strains N2, CB4856, CB4852, DL238, ECA369, EG4348, JU1088, JU1581, KR314, and QX1211 were reared under standard conditions (as above), bleached to isolate embryos, and grown to reproductive maturity. RNA was extracted with TRIzol (Invitrogen #15596026) and RNeasy columns (Qiagen #74104), following (*He, 2011*). RNA was collected at two timepoints, early and middle reproductive maturity (68±2hrs and 90±2hrs after bleaching, respectively). RNA sample concentrations were quantified and standardized using a Nanodrop (Thermo Scientific), and cDNA was synthesized using the ProtoScript II First Stand cDNA Synthesis Kit (NEB #E6560S). The experiment was replicated as follows: from each experimental condition, we collected two RNA samples, for two biological replicates; within the plate, each reaction was duplicated, for two technical replicates; and we conducted the entire experiment twice.

Droplet digital PCR was carried out with the Bio-Rad QX200 system following the manufacturer’s protocol, and results were obtained using the QuantaSoft software (Bio-Rad), via automatic thresholding followed by manual confirmation of droplet selection. All samples produced >8000 droplets and results from all samples were retained. Concentration, given by number of copies per μL, was modeled with a quasipoisson error structure using the *glm()* function in R. As *ppw-1* was detected at an order of magnitude higher than *sago-2*, we analyzed the two genes separately. By model selection, we identified the minimal model that best described the observed differences in concentration For the *ppw-1* analysis, we dropped run date from the model, as it was not significant; for *sago-2*, run date contributed <1% to the total observed deviance (Table S2), but was nevertheless significant, so it was retained. The final models were: Concentration ∼ Strain*DevStage/BiolRep for *ppw-1*, and Concentration ∼ RunDate + Strain*DevStage/BiolRep for *sago-2*. To determine which strains differed in *ppw-1* or *sago-2* levels, we performed pairwise contrasts among strains using the *TukeyHSD()* function and a family-wise confidence level of 95% (only a subset of comparisons are reported in the text). To determine which strains showed differences in concentration according to developmental stage, we performed pairwise contrasts using the *lsmeans()* function in the package *lsmeans*, using a confidence level of 95% following a Bonferroni correction for multiple tests.

### Genotype and sequence analysis

To evaluate population-level allelic variation at known RNAi genes, we queried the *C. elegans* Natural Diversity Resource (https://elegansvariation.org), which provides genotype data for 403 wild isotypes from short read sequence data mapped to the N2 reference genome (*Cook et al., 2017*). Specifically, we downloaded the VCF (WI.20200815.hard-filter.isotype.vcf) and used the R package *VariantAnnotation* (*Obenchain et al., 2014*) to extract information about mutations and mapping coverage, and the package *PopGenome* (*Pfeifer et al., 2014*) to estimate nucleotide diversity, at each gene. Haplotype networks for individual genes were determined using the R package *pegas* (*Paradis, 2010*).

For a subset of strains, we verified and/or supplemented the genotype data with de novo-assembled genome data and long read data. Genomic DNA of strains AB2, EG4347, EG4348, JU1088, JU1171, PB306, PX174, QX1211 and QX1216 was prepared using standard phenol/chloroform extraction and ethanol precipitation. Samples were cleaned with DNA Clean & Concentrator columns (Zymo Research #D4004) and libraries were prepared using NEBNext Ultra II FS DNA Library Prep Kit for Illumina (New England Biolabs #E7805) and Multiplex Oligos for Illumina (NEB #E7500), with customized fragmentation and purification steps to enrich for desired sizes. A final DNA size selection targeting 650bp±50bp was performed using BluePippin (Sage Science). The libraries were sequenced on a HiSeq 2500 (Illumina) on Rapid Run Mode (paired-end 2x250bp) in the Molecular Evolution Core at the Georgia Institute of Technology. Raw data were trimmed using Cutadapt (v1.18) (*Martin, 2011*) and quality control was performed with FastQC (*Andrews, 2021*). Reads were then assembled into contigs with DISCOVAR *de novo* (v52488) (Broad Institute) using default parameters. Separately, QX1211 and JU1088 genomic DNA samples were snap frozen with liquid nitrogen and sent to the Georgia Genomics and Bioinformatics Core (GGBC) at the University of Georgia. Quality was assessed by Qubit (Invitrogen) and NanoDrop (Thermo Scientific), molecular weight distribution was assessed by fragment analysis, and sizes >15kb were selected by BluePippin (Sage Science). Each sample was sequenced on a single SMRT Cell on the PacBio Sequel I platform (Pacific Biosciences). Genome assembly was performed by the GGBC using Canu (v1.7) (*Koren et al., 2017*).

### Computing

Unless otherwise specified, all analyses were performed in R (*R Core Team, 2021*) and figures were generated with the packages *ggplot2* (*Wickham, 2016*) and *ggpubr* (*Kassambara, 2020*). Computationally intensive jobs, including read mapping and genome assembly, were performed on PACE, the high performance computing platform at the Georgia Institute of Technology.

## ACKNOWLEDGEMENTS

We wish to acknowledge the core facilities at the Parker H. Petit Institute for Bioengineering and Bioscience at the Georgia Institute of Technology for the use of their shared equipment, services and expertise. Specifically, we are grateful to Aaron Lifland at the Optimal Microscopy Core for training in confocal imaging, and Anton Bryksin, Naima Djeddar, and Shweta Biliya at the Molecular Evolution Core for collaboration on the sequencing projects. We also thank the Georgia Genomics and Bioinformatics Core at the University of Georgia for PacBio sequencing and genome assembly. We thank Miyeko Mana for the gift of the *par-4* RNAi clone, and Lijiang Long and Patrick McGrath for strain PTM377. Some strains were provided by the CGC, which is funded by NIH Office of Research Infrastructure Programs (P40 OD010440). This research was supported in part through research cyberinfrastructure resources and services provided by the Partnership for an Advanced Computing Environment (PACE) at the Georgia Institute of Technology. This research was funded by NIH grant R35 GM119744 to A.B.P, NSF grant MCB1518314 to D.A.P., and NSF fellowship 2109666 to A.D.B.

## SUPPLEMENT

**Figure S1.**
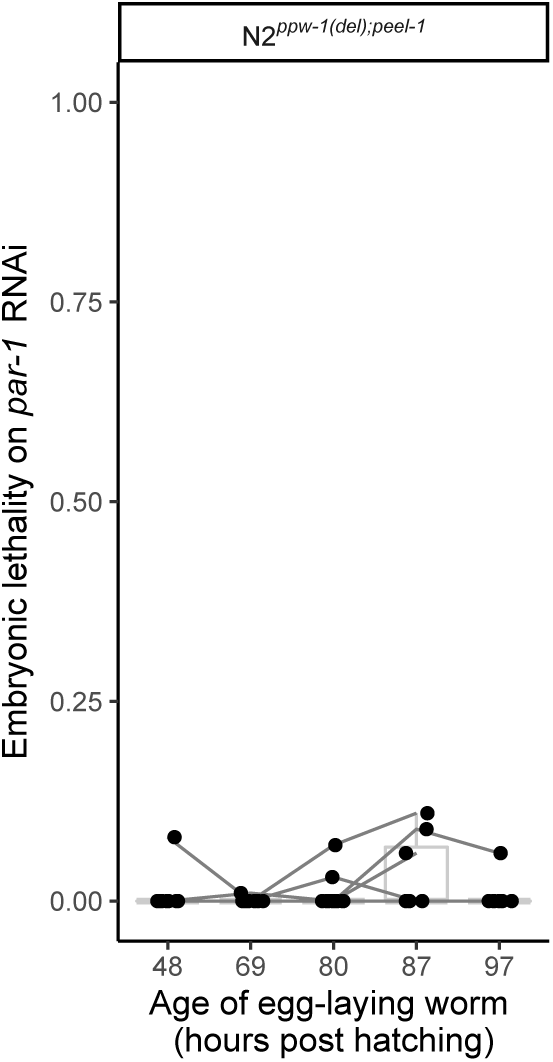
Embryonic lethality following RNAi against *par-1* in the *ppw-1;peel-2* mutant.

**Figure S2.**
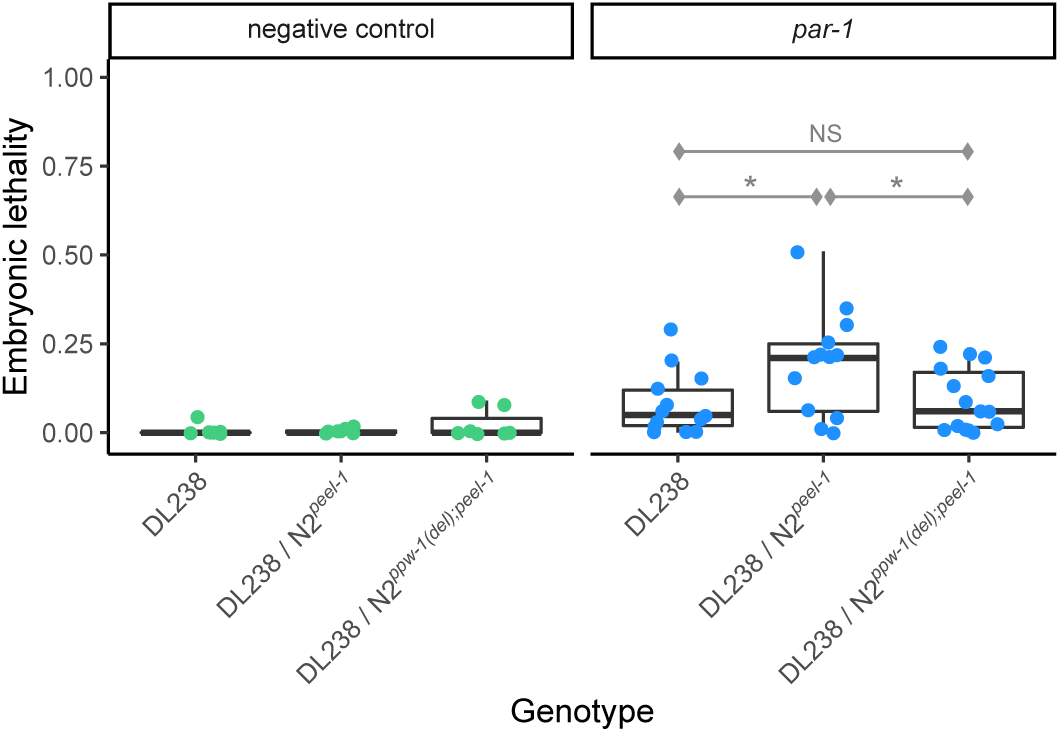
Complementation test for DL238 with pooled hermaphrodites. DL238 was crossed to the RNAi-sensitive laboratory strain N2, with and without a deletion allele at *ppw-1*. The embryos scored here were laid by pooled hermaphrodites on replicate plates. Significance level (Tukey’s contrasts): p<0.05 (*).

**Figure S3.**
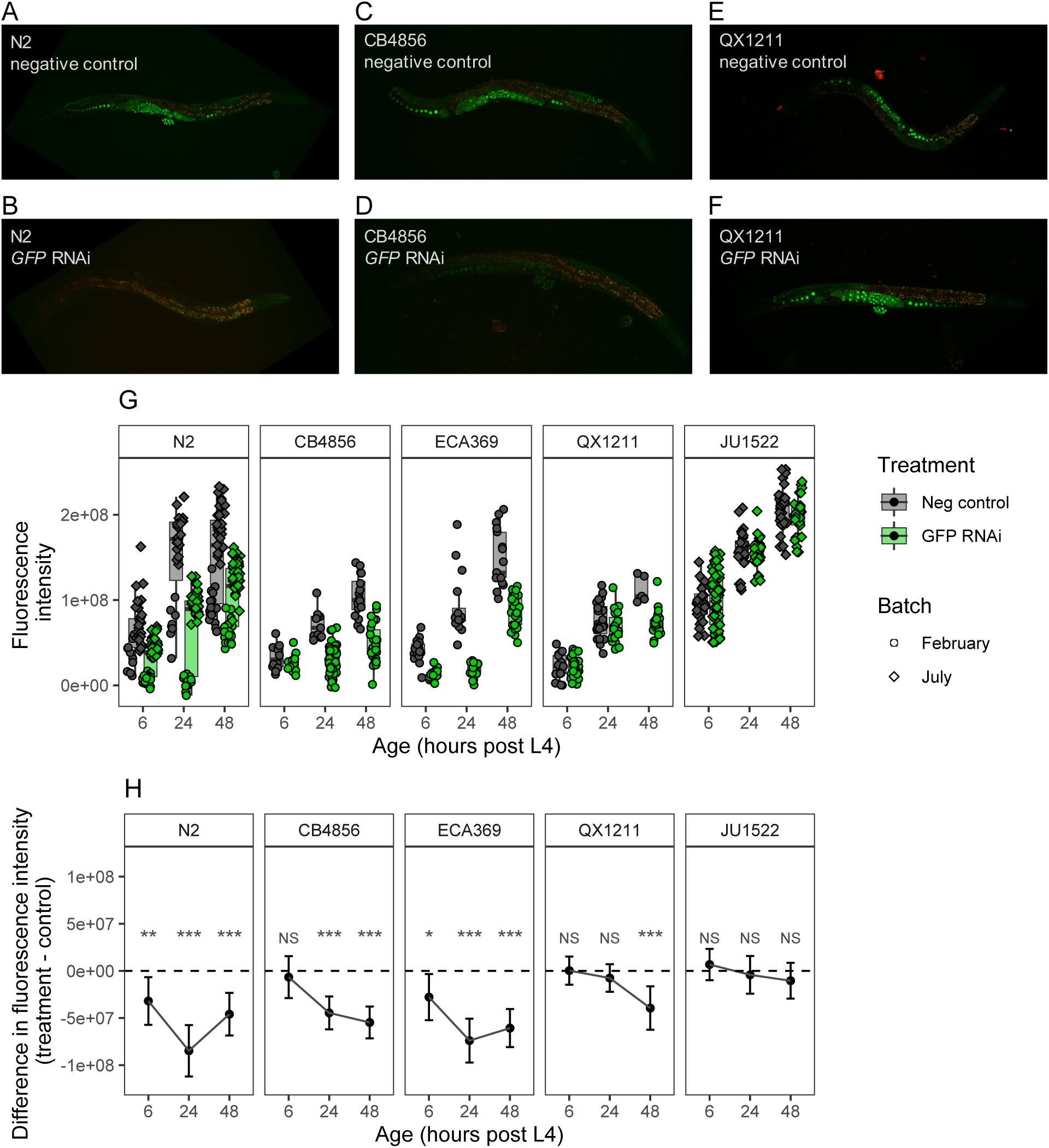
RNAi against germline-expressed GFP in wild-type strains. (A) - (F) Representative images of control and treated worms, all at 24hrs post L4 stage, are shown for N2, CB4856 and QX121. Green indicates GFP fluorescence; the red channel is overlaid on these images to show autofluorescence. (G) Whole-worm fluorescence intensity for worms imaged at 6, 24, and 48hrs post L4 stage; each point represents fluorescence measured for a single individual. (H) The difference in fluorescence intensity between treated and untreated samples; error bars represent standard error. Significance levels (Tukey’s contrasts): p<0.001 (***), p<0.01 (**), p<0.05 (*).

**Figure S4.**
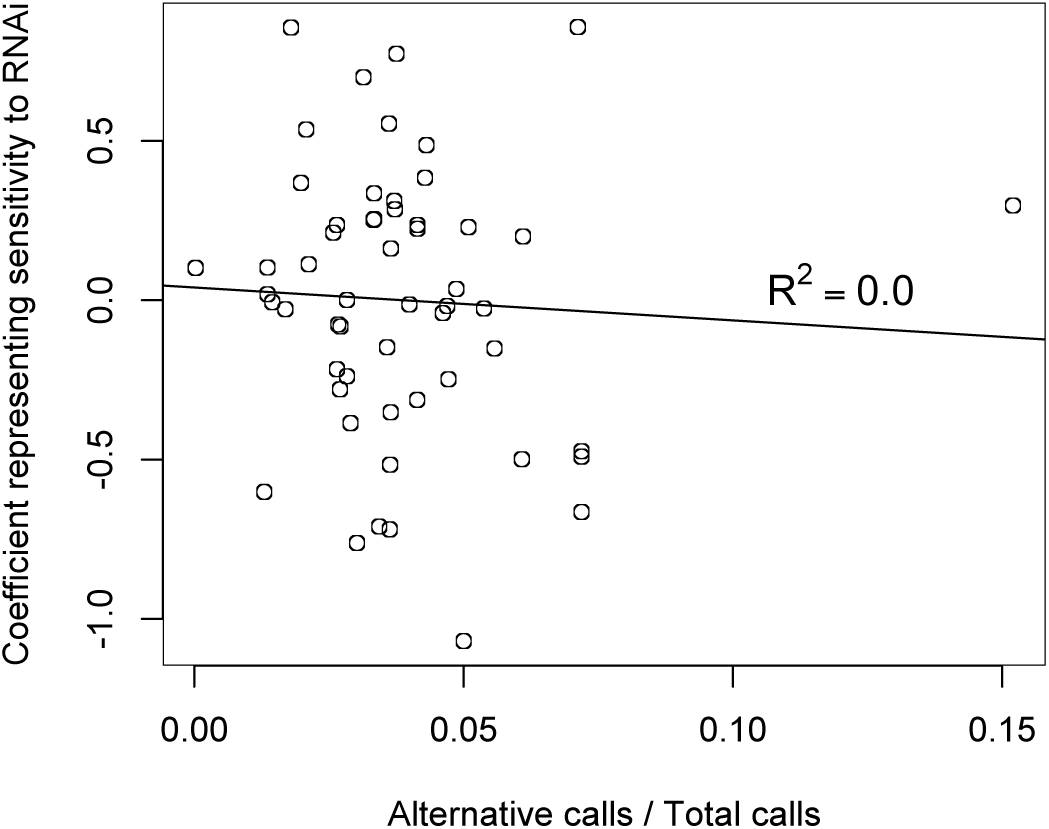
RNAi sensitivity for 55 wild isolates versus genetic distance from reference strain N2. RNAi data is from (*Paaby et al., 2015*), which measured embryonic lethality following RNAi by feeding against 29 individual maternal-effect targets. The y-axis plots the coefficients associated with the strain term in the full statistical model, and represents the strain-specific variation in embryonic lethality associated with RNAi responsiveness; increasingly positive values indicate weaker germline RNAi (*Paaby et al., 2015*). The x-axis gives the proportion of alternate genotype calls, out of the total number of calls, for each strain in the CeNDR database (*Cook et al., 2017*).

**Figure S5.**
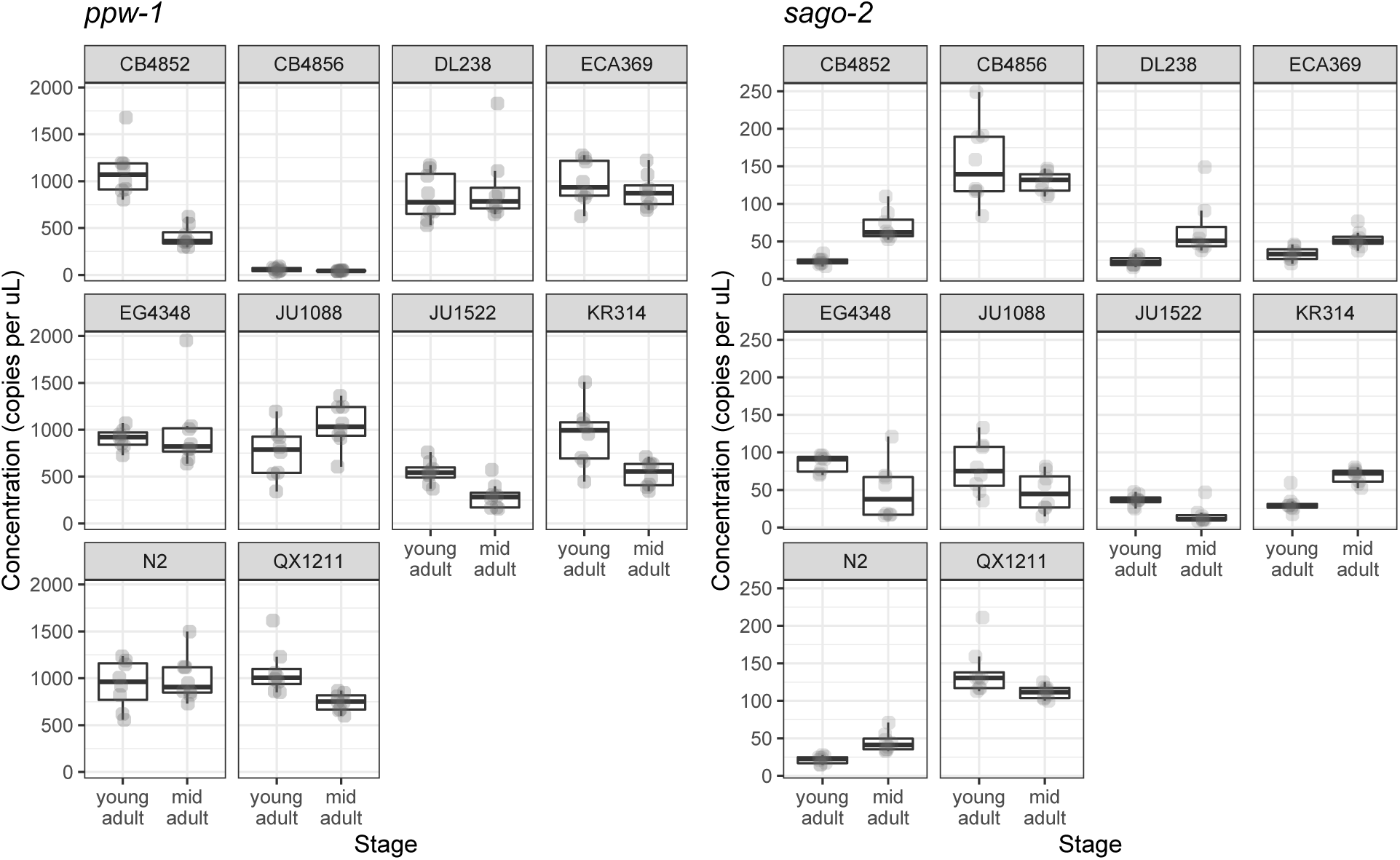
Droplet digital PCR results for *ppw-1* and *sago-2* for all ten tested strains.

**Figure S6.**
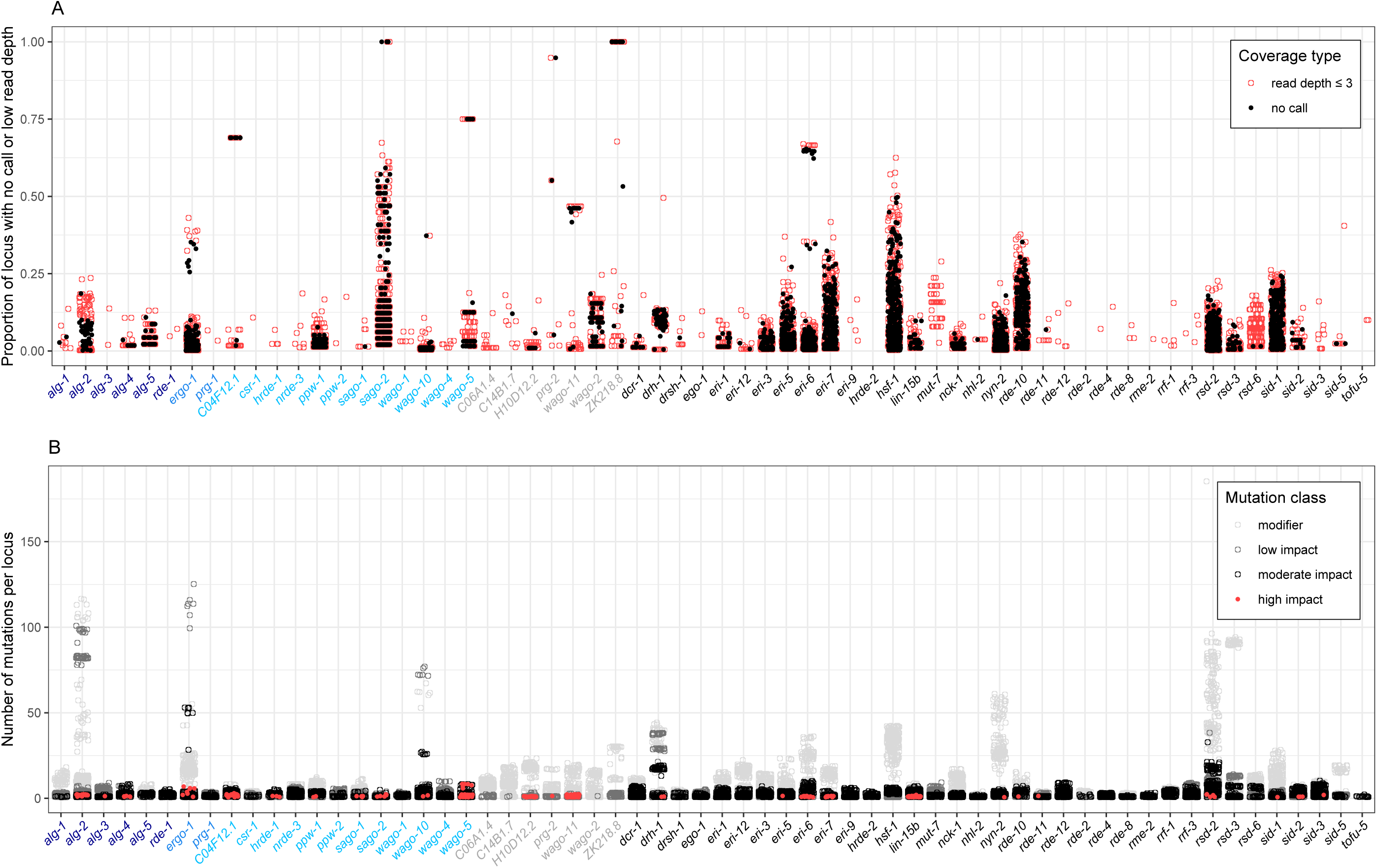
Population-level sequence variation for 62 RNAi genes. (A) mapping coverage and (B) mutational variation is shown for the 403 strain isotypes from CeNDR. Gene names are color coded by classification: AGO Argonaute (dark blue), PIWI Argonaute (blue), WAGO Argonaute (light blue), Argonaute pseudogene (grey), and other RNAi factor (black). Each point corresponds to an individual strain; zeros (no observation of poor coverage or mutation relative to the reference) are not plotted.

**Table S1.** Statistical estimates for changes in smFISH transcript abundance. The interaction term represents the changes in slope (Figure 2) between treated and untreated samples.

| | | Variance explained ($\omega^2$) | p-value (ANCOVA model comparison) |
| --- | --- | --- | --- |
| N2 | Treatment | 0.586 | <0.001 |
|  | Embryo stage | 0.248 | <0.001 |
|  | Interaction | 0.059 | <0.001 |
| CB4856 | Treatment | 0.010 | 0.043 |
|  | Embryo stage | 0.581 | <0.001 |
|  | Interaction | 0.017 | 0.012 |
| QX1211 | Treatment | 0.014 | 0.037 |
|  | Embryo stage | 0.208 | <0.001 |
|  | Interaction | NA | NS |

**Table S2.** Statistical results for the ddPCR analysis. The model used for each gene is described in the Methods.

|  | Df | Deviance | Resid. Df | Resid. Dev |
| --- | --- | --- | --- | --- |
| NULL |  |  | 158 | 37942 |
| Strain | 9 | 24433.1 | 149 | 13509 |
| DevStage | 1 | 910.3 | 148 | 12599 |
| Strain:DevStage | 9 | 4271.9 | 139 | 8327 |
| Strain:DevStage:BiolRepl | 20 | 2327.6 | 119 | 5999 |

*sago-2*
|  | Df | Deviance | Resid. DF | Resid. Dev |
| --- | --- | --- | --- | --- |
| NULL |  |  | 158 | 4722.0 |
| RunDate | 1 | 29.38 | 157 | 4692.6 |
| Strain | 9 | 2932.73 | 148 | 1759.9 |
| DevStage | 1 | 5.87 | 147 | 1754.0 |
| Strain:DevStage | 9 | 825.20 | 138 | 928.8 |
| Strain:DevStage:BiolRepl | 20 | 436.26 | 118 | 492.6 |

**Table S3.** Candidate genes for weak germline RNAi in the seven strains tested. Only genes with relevant genotypes in at least one strain are displayed, out of 62 RNAi genes queried. “High impact” indicates disruptive mutations like frameshifts or stop-gains. “Poor mapping coverage” indicates a read depth of 3 or less. “Functional diverged allele” indicates at least 1% nucleotide divergence from the reference genome, including at least five moderate mutations, such as amino acid substitutions, and no high impact mutations.

| Class | Gene | One or more high impact mutation | Poor mapping coverage at 10% or more of locus | Functional diverged allele |
| --- | --- | --- | --- | --- |
| Argonaute (AGO) | *alg-4* |  |  | QX1211 |
| Argonaute (PIWI) | *ergo-1* | ECA369 | ECA369 |  |
| Argonaute (WAGO) | *hrde-1* |  |  | QX1211 |
|  | *nrde-3* |  |  | QX1211 |
|  | *ppw-1* | CB4856 |  |  |
|  | *sago-2* | DL238 | DL238, ECA369, JU1522, KR314, QX1211 |  |
|  | *wago-10* |  |  | QX1211 |
| Argonaute (pseudogene) | *wago-11* | CB4852, KR314, QX1211 | CB4856 |  |
|  | *wago-2* |  | QX1211 |  |
|  | *ZK218.8* |  | DL238 |  |
| Other RNAi factor | *dcr-1* |  |  | QX1211 |
|  | *drh-1* |  |  | QX1211 |
|  | *eri-1* |  |  | QX1211 |
|  | *eri-5* |  | JU1522 |  |
|  | *eri-6* |  |  | QX1211 |
|  | *eri-7* | CB4856, DL238, ECA369 | DL238, JU1522, KR314 |  |
|  | *eri-9* |  |  | QX1211 |
|  | *lin-15b* | QX1211 |  | ECA369 |
|  | *nyn-2* |  | DL238 | QX1211 |
|  | *rde-10* |  | DL238, ECA369, JU1522, KR314, QX1211 |  |
|  | *rde-12* |  |  | QX1211 |
|  | *rsd-2* |  | JU1522, QX1211 | CB4856, DL238, ECA369, QX1211 |
|  | *sid-1* |  | DL238, ECA369, JU1522, QX1211 |  |
|  | *sid-2* |  |  | QX1211 |
|  | *sid-3* |  |  | QX1211 |
|  | *hsf-1* |  | DL238, ECA369, JU1522, KR314 | ECA369, KR314 |

**Table S4.**
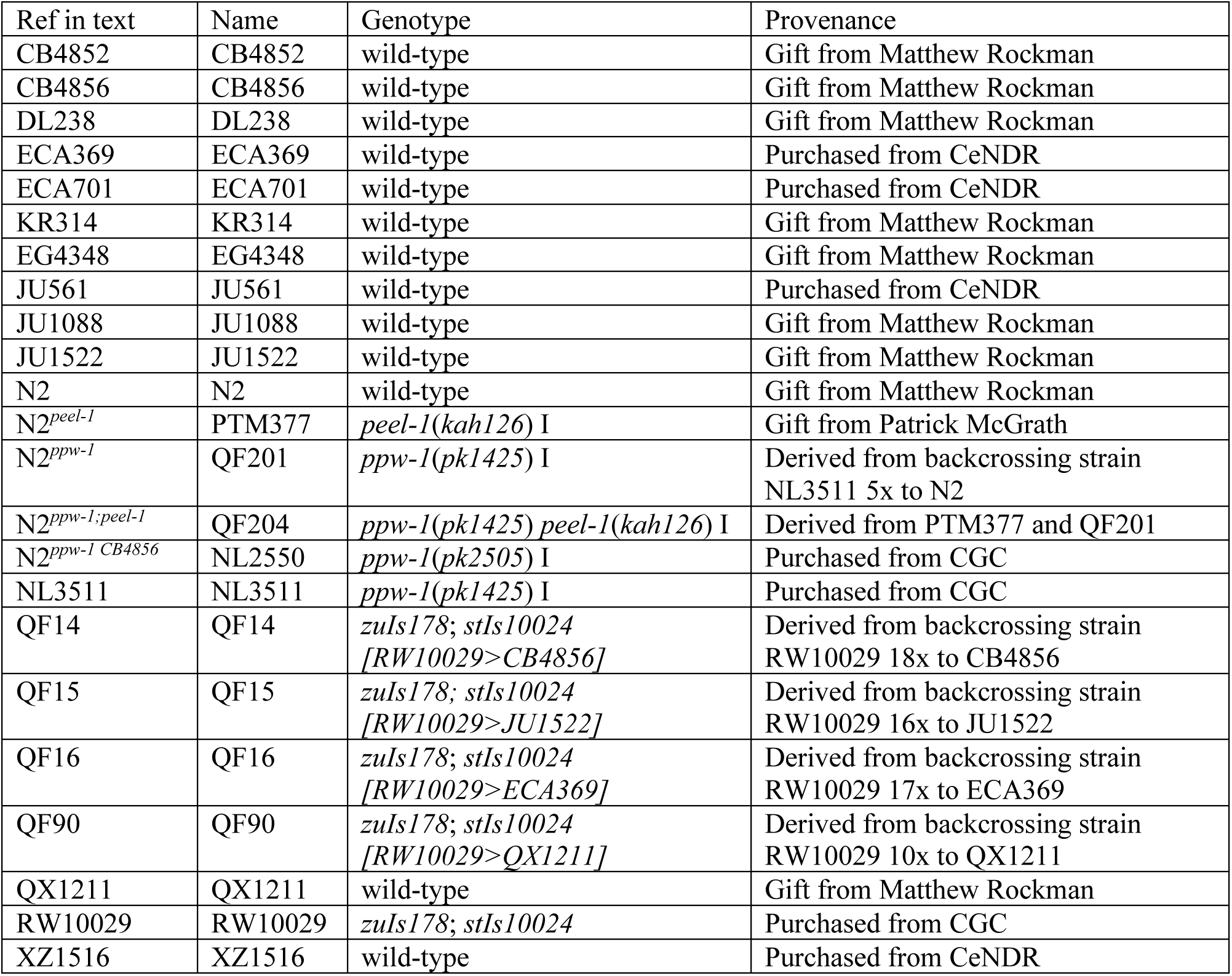
Strains used in this study.

| Ref in text | Name | Genotype | Provenance |
| --- | --- | --- | --- |
| CB4852 | CB4852 | wild-type | Gift from Matthew Rockman |
| CB4856 | CB4856 | wild-type | Gift from Matthew Rockman |
| DL238 | DL238 | wild-type | Gift from Matthew Rockman |
| ECA369 | ECA369 | wild-type | Purchased from CeNDR |
| ECA701 | ECA701 | wild-type | Purchased from CeNDR |
| KR314 | KR314 | wild-type | Gift from Matthew Rockman |
| EG4348 | EG4348 | wild-type | Gift from Matthew Rockman |
| JU561 | JU561 | wild-type | Purchased from CeNDR |
| JU1088 | JU1088 | wild-type | Gift from Matthew Rockman |
| JU1522 | JU1522 | wild-type | Gift from Matthew Rockman |
| N2 | N2 | wild-type | Gift from Matthew Rockman |
| N2<sup>peel-1</sup> | PTM377 | *peel-1(kah126)* I | Gift from Patrick McGrath |
| N2<sup>ppw-1</sup> | QF201 | *ppw-1(pk1425)* I | Derived from backcrossing strain NL3511 5x to N2 |
| N2<sup>ppw-1;peel-1</sup> | QF204 | *ppw-1(pk1425) peel-1(kah126)* I | Derived from PTM377 and QF201 |
| N2<sup>ppw-1 CB4856</sup> | NL2550 | *ppw-1(pk2505)* I | Purchased from CGC |
| NL3511 | NL3511 | *ppw-1(pk1425)* I | Purchased from CGC |
| QF14 | QF14 | *zuIs178; stIs10024*<br>[RW10029>CB4856] | Derived from backcrossing strain RW10029 18x to CB4856 |
| QF15 | QF15 | *zuIs178; stIs10024*<br>[RW10029>JU1522] | Derived from backcrossing strain RW10029 16x to JU1522 |
| QF16 | QF16 | *zuIs178; stIs10024*<br>[RW10029>ECA369] | Derived from backcrossing strain RW10029 17x to ECA369 |
| QF90 | QF90 | *zuIs178; stIs10024*<br>[RW10029>QX1211] | Derived from backcrossing strain RW10029 10x to QX1211 |
| QX1211 | QX1211 | wild-type | Gift from Matthew Rockman |
| RW10029 | RW10029 | *zuIs178; stIs10024* | Purchased from CGC |
| XZ1516 | XZ1516 | wild-type | Purchased from CeNDR |

### File S1. Statistical details of smFISH in early stage embryos

In early stage embryos (up to four cells), N2 showed a significant reduction in *par-1* transcript abundance after *par-1* RNAi (t=-16.34, df=32, p<0.001), while CB4856 and QX1211 did not (t=0.41, df=56.31, p=0.684 and t=1.15, df=95.03, p=0.255, respectively) (Figure 2D). However, for CB4856, the variance in abundance was slightly higher for treated embryos than for control embryos (F=1.94, df=32,40, p=0.048), which may represent the onset of an RNAi response. In QX1211, the variance of *par-1* treated embryos was substantially higher (F=2.94, df=100,32, p<0.001) and abundance was bimodally distributed, with low-abundance embryos at levels similar to N2; *par-4* treated embryos showed a slight reduction in average abundance and substantial increase in variance (t=-2.64, df=24.20, p=0.014 and F=5.50, df=20,36, p<0.001, respectively) (Figure 2D).

### File S2. Details of genetic incompatibilities

*C. elegans* carries two known genetic incompatibilities: the paternally-delivered toxin *peel-1*, which is rescued by the zygotically-expressed antidote *zeel-1* (*Seidel et al., 2008,* 2011), and the maternally-delivered toxin *sup-35*, which is rescued by the zygotically-expressed antidote *pha-1 (Ben-David et al., 2017)*. In both instances, embryos that cytologically inherit the toxin but do not inherit the genotype to express the antidote will die. N2 carries both toxin-antidote complexes but several of our other wild isolates do not. Since our goal was to use embryonic lethality to measure the RNAi response, lethality arising from these genetic incompatibilities had the potential to confound our results.

To control for the *zeel-1;peel-1* incompatibility, we generated a strain derived from N2 with both the *ppw-1* deletion and an allele of *peel-1* (*kah126*) that disables the toxin by a frameshift insertion in the second exon (N2*^ppw-1(del);peel-1^*). This allele eliminates embryonic lethality that would otherwise arise in the F2 generation, from F1 heterozygotes derived from incompatible strains. The presence of this *peel-1* allele had no effect on our measured RNAi response in either sensitive or resistant backgrounds (Figure 3A, Figure S1), so we used it in all comparisons requiring the N2 background.

We did not control for the *sup-35;pha-1* incompatibility. We anticipated toxin-associated embryonic lethality to occur from crosses initiated between N2 and two wild isolates without the active *sup-35;pha-1* complex: DL238 and QX1211. For DL238, we observed only very weak penetrance of this effect (Figure 3C). For QX1211, we observed clear toxin-associated lethality, as embryos derived from the N2 × QX1211 cross showed lethality on the control condition (Figure 3D). However, we were still able to infer a distinct *ppw-1*-associated response in this assay, as lethality from the QX1211/N2*^peel-1^* genotype matched that of the control and lethality from QX1211/N2*^ppw-1(del);peel-1^* was significantly higher (Figure 3D).

## Notes

### Competing Interest Statement

The authors have declared no competing interest.

